# The PfAP2-G2 transcription factor is a critical regulator of gametocyte maturation

**DOI:** 10.1101/2020.10.27.355685

**Authors:** Suprita Singh, Joana M. Santos, Lindsey M. Orchard, Naomi Yamada, Riëtte van Biljon, Heather J. Painter, Shaun Mahony, Manuel Llinás

**Affiliations:** Department of Biochemistry and Molecular Biology, The Pennsylvania State University, University Park, PA, USA 16802, Huck Center for Malaria Research, The Pennsylvania State University, University Park, PA, USA 16802; Center for Eukaryotic Gene Regulation, Department of Biochemistry & Molecular Biology, The Pennsylvania State University, University Park, PA, USA 16802; Department of Chemistry, The Pennsylvania State University, University Park, PA, USA 16802

## Abstract

Differentiation from asexual blood stages to sexual gametocytes is required for transmission of malaria parasites from the human to the mosquito host. Preventing gametocyte commitment and development would block parasite transmission, but the underlying molecular mechanisms behind these processes remain poorly understood. Here, we report that the ApiAP2 transcription factor, PfAP2-G2 (PF3D7_1408200) plays a critical role in the maturation of *Plasmodium falciparum* gametocytes. PfAP2-G2 binds to the promoters of a wide array of genes that are expressed at many stages of the parasite life cycle. Interestingly, we also find binding of PfAP2-G2 within the gene body of almost 3000 genes, which strongly correlates with the location of H3K36me3 and several other histone modifications as well as Heterochromatin Protein 1 (HP1), suggesting that occupancy of PfAP2-G2 in gene bodies may serve as an alternative regulatory mechanism. Disruption of *pfap2-g2* does not impact asexual development, parasite multiplication rate, or commitment to sexual development but the majority of sexual parasites are unable to mature beyond stage III gametocytes. The absence of *pfap2-g2* leads to overexpression of 28% of the genes bound by PfAP2-G2 and none of the PfAP2-g2 bound are downregulated, suggesting that it is a repressor. We also find that PfAP2-G2 interacts with chromatin remodeling proteins, a microrchidia (MORC) protein, and another ApiAP2 protein (PF3D7_1139300). Overall our data demonstrate that PfAP2-G2 is an important transcription factor that establishes an essential gametocyte maturation program in association with other chromatin-related proteins.

## Introduction

Malaria is a life-threatening disease that continues to impact the lives of millions of people worldwide. According to the latest WHO estimates, there were 228 million cases of malaria in 2018, resulting in 405,000 deaths (World Malaria Report, WHO, 2019). Malaria is caused by unicellular protozoan parasites belonging to the genus *Plasmodium*. Of the six species that infect humans, *Plasmodium falciparum* has the highest mortality rate (Weiss et al. 2019; WHO 2019). *P. falciparum* exhibits a complex life cycle with asexual and sexual erythrocytic phases in the human host, followed by development in *Anopheles* mosquitoes which transmit the parasite back to a new host, initiating the pre-erythrocytic liver stage of infection. Although the cyclic 48-hour asexual blood-stage is responsible for symptomatic disease and severe malaria, this form of the parasite cannot be transmitted. Rather, in each round of replication, a fraction of the parasites <10% exit the asexual pathway and undergo sexual differentiation to form male and female gametocytes, which are transmission competent (Bruce et al. 1989). Inhibition of gametocyte development is of great interest, because it would prevent malaria parasite transmission, which is one of the major goals in the effort to achieve disease eradication (Rabinovich et al. 2017).

Gametocyte development in *P. falciparum* is a 9–12 day process, which is substantially longer than that of other human infecting *Plasmodium* species such as *P. vivax* and the rodent malaria parasites *P. berghei* (26–30 hours) and *P. yoelii* (36 hours) (Gautret and Motard, 1999; Liu, Miao, & Cui, 2011; Armistead et al., 2018). *P. falciparum* gametocyte development is divided into five (stage I to V) morphologically distinct stages (Sinden 1982). The earliest phase of gametocyte development, stage Ia, occurs around 24 to 30 hours post-invasion (hpi) and is morphologically indistinguishable from the young trophozoite stage. However, there are several well-defined markers of early stage I gametocytes such as Pfs16 and Pfg27 and PfGEXP-5 (Alano, Premawansa, Bruce, & Carter, 1991; Marian C. Bruce, Carter, Nakamura, Aikawa, & Carter, 1994; Silvestrini et al., 2010; Poran et al., 2017; Josling et al., 2020; Llorà-Batlle et al., 2020). In the human host, stage Ib to stage IV sexually-developing parasites are sequestered in deep tissues like the bone marrow and only stage V gametocytes freely circulate in the blood and can be picked up by mosquitoes (Aguilar et al. 2014; Joice et al. 2014; Venugopal et al. 2020)

Regulation of *Plasmodium* development is driven by stage-specific transcription factors (TFs), such as the well-studied Apicomplexan AP2 (ApiAP2) family of DNA binding proteins. ApiAP2 proteins are found among all members of the phylum, and each ApiAP2 protein contains between one to three Apetala2 (AP2) DNA binding domains (Balaji et al. 2005). In *P. falciparum* there are 27 members of the ApiAP2 protein family. ApiAP2 proteins have been shown to control all developmental transitions in *Plasmodium* (Jeninga, Quinn, and Petter 2019; Modrzynska et al. 2017; Zhang et al. 2017)

A master regulator of sexual (gametocyte) commitment, AP2-G, has been identified in both *P. falciparum* and *Plasmodium berghei*, a rodent malaria parasite (Kafsack et al. 2014; Sinha et al. 2014). Expression levels of PfAP2-G (PF3D7_1222600) strongly correlate with the formation of gametocytes, and targeted disruption of the *pfap2-g* locus results in a complete block in gametocytogenesis and downregulation of many gametocyte-associated genes (Kafsack et al. 2014). Another regulator of *Plasmodium* gametocyte development is AP2-G3. Disruption of *pyap2-g3* (PY17X_1417400) in *P. yoelii* resulted in significant reduction in the numbers of male and female gametocytes, day 8 oocysts, and sporozoites, but it did not affect the asexual growth of the parasites (Zhang et al. 2017). The *P. berghei* orthologue, PBANKA_1415700, has also been knocked out and shown to be female-specific and essential for female gametocyte development, and was thus renamed PbAP2-FG (Yuda et al. 2019). The *P. falciparum* orthologue, PF3D7_1317200, was also found to be associated with gametocytogenesis (Ikadai et al. 2013), although it has not been extensively characterized.

AP2-G2 has been shown to play a role in gametocytogenesis in both *P. berghei* and *P. yoelii* (Sinha et al., 2014; Yuda, Iwanaga, Kaneko, & Kato, 2015; Modrzynska et al., 2017). Knockout of *pbap2-g2* does not inhibit sexual stage conversion but rather results in the nearly complete loss of gametocyte maturation and a block in transmission to mosquitoes (Yuda et al. 2015). Yuda *et al*. reported that the *P. berghei* transcription factor is bound to roughly 1,500 genes, or slightly over 1/3 of the genome during asexual development, and a number of these genes were up-regulated by more than two-fold in *pbap2-g2* knockout lines. In another study, disruption of *pbap2-g2* also caused premature expression of liver stage and sporozoite stage genes during asexual development in red blood cells (Modrzynska et al. 2017). Moreover, a *P. berghei* liver-stage (LS) transcriptome of *P. berghei* reported strong upregulation of *pbap2-g2* in early LS that is negatively correlated with the expression of liver stage-specific transcripts (Derbyshire 2019). Together, these findings suggest that AP2-G2 acts as a repressor in both the asexual and sexual stages and during early liver-stage infection in *P. berghei*. Accordingly, *P. yoelii* parasites lacking the *pyap2-g2* gene had greatly reduced numbers of gametocytes and oocysts (Zhang et al. 2017). A recent genome-wide knockout screen suggests that AP2-G2 is also not essential for blood-stage development in *P. falciparum* (Zhang et al. 2018), and may have a role in a later stage of development.

In this study we explore the function of AP2-G2 in *P. falciparum* during parasite blood-stage development. The period of gametocyte maturation for rodent malaria parasites is shorter than in *P. falciparum*, making it difficult to discern the developmental phenotypes associated with *ap2-g2* knockouts. Therefore, disrupting this gene in *P. falciparum* is of interest due to the longer maturation period. Furthermore, two studies have reported differences in asexual blood-stage development resulting from *pbap2-g2* deletion. On one hand, Modrzynska *et al*. observed reduced growth and premature expression of liver- and sporozoite-stage genes (Modrzynska et al. 2017). Conversely, Yuda *et al*. reported that *pbap2-g2* KO parasites proliferate normally in blood (Yuda et al. 2015). Another discrepancy regards the impact of the KO on the sex ratio. Sinha *et al*. reported a difference in the ratio of male to female gametocytes was disrupted in *pbap2-g2* KO lines (Sinha et al. 2014) while Yuda et al. observed no such differences (Yuda et al. 2015).

We also find that the *P. falciparum orthologue* of AP2-G2 also plays a critical role in the maturation of gametocytes. Disruption of *pfap2-g2* does not impact asexual development, parasite multiplication rate, or commitment to sexual development, but parasites are unable to develop normally beyond stage III. Using transcriptomic analysis and ChIP-seq we have identified a number of candidate genes that are bound and regulated by PfAP2-G2. We also identify interacting partners using protein immunoprecipitation followed by mass spectrometry. Overall our work suggests that PfAP2-G2 is a transcriptional repressor that likely recruits additional transcription factors and chromatin remodeling machinery to the genome to control gene expression. PfAP2-G2 plays a critical role in the regulation of the development of malaria parasites as they transition from asexual to sexual parasites, allowing for proper gametocyte maturation.

## Results

### PfAP2-G2 is expressed during the trophozoite and schizont stages of asexual development

The *P. falciparum* orthologue of AP2-G2, PF3D7_1408200, encodes a 189kDa protein that contains a single AP2 DNA binding domain (**Figure 1A**). *pfap2-g2* is maximally transcribed at the ring and early trophozoite stages (Bozdech et al., 2003; Painter et al., 2018) and proteomics data indicates protein expression at the trophozoite and schizont stages (Oehring et al. 2012). To precisely determine the specific stage of expression of PfAP2-G2and its subcellular localization, we tagged the gene at the C-terminus with GFP (**Supplementary Fig. 1**). Using live-cell fluorescence microscopy, we imaged a highly synchronized PfAP2-G2::GFP parasite population for one complete asexual replication cycle (48-hours), every 7 h beginning with the newly invaded ring stage (5–7 hpi) revealed that PfAP2-G2 was localized to the nucleus, as expected, and was expressed from the early trophozoite to the late schizont stages (**Figure 1B**). Further confirming the nuclear localization of PfAP2-G2, nuclear fractionation of trophozoite stage parasites (~30 hpi) showed that full length PfAP2-G2::GFP (~250 kDa) was only detected in the nuclear fraction of the parasite lysates (**Figure 1C**). We also detected other smaller sized peptides indicating that PfAP2-G2 may be proteolytically cleaved or is unstable in parasite lysates.

**Figure 1.**
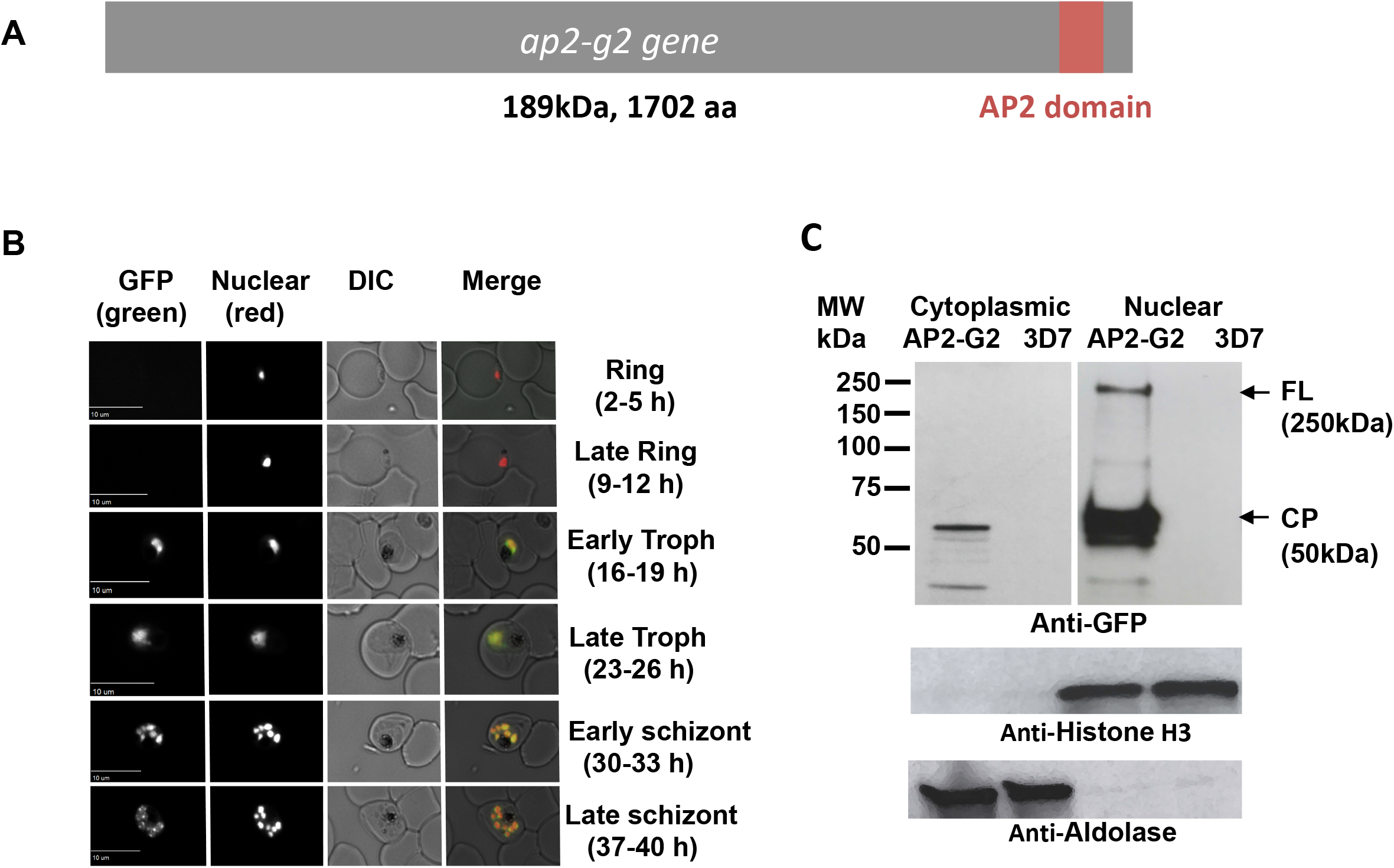
PfAP2-G2 is expressed from around 16-19 hpi in asexual stage parasite. (A) Structure of PfAP2-G2 protein which contains single AP2 DNA-binding domain. (B) Live fluorescence microscopy performed on a highly synchronized population of AP2-G2::GFP parasites every 7 h of the asexual life cycle showed that PfAP2-G2::GFP colocalizes with the nucleus in both the trophozoite and schizont stages. DRAQ-5 was used as the nuclear stain. (C) Nuclear and cytoplasmic fractions from the GFP tagged PfAP2-G2 synchronized trophozoite stage parasites were subjected to western blot analysis using anti-GFP antibodies (1:1000). Full length (FL) GFP-tagged PfAP2-G2 (~250 kDa) was detected only in the nuclear fraction of the parasite lysate whereas cleavage product (CP, ~50kDa) was in both fractions. WT parasites were used as a negative control. Anti-histone H3 (1:3000) and anti-aldolase (1:1000) were used as loading controls for the nuclear and cytoplasmic fractions, respectively.

### PfAP2-G2 is not required for proliferation during the asexual stages of the life cycle

To determine the role *of pfap2-g2*, we generated a genetic disruption (KO) line using selection linked integration (SLI) (Birnbaum et al. 2017) to truncate the *pfap2-g2* coding sequence (**Supplementary Fig. 2A**). As expected, *pfap2-g2* KO parasites were readily obtained. Live-cell fluorescence imaging of the transgenic parasites showed diffuse cytoplasmic staining suggesting that the remaining non AP2 domain-containing protein fragment no longer localized to the nucleus (**Supplementary Fig. 2B**). We next determined if truncation of PfAP2-G2 impacted asexual stage parasite growth or multiplication rate. Using a SYBR green growth assay (Vossen et al. 2010), we find that *pfap2-g2* KO parasites develop similarly to that of WT (**Figure 2A**), and have similar multiplication rates (**Figure 2B**). Therefore, although PfAP2-G2 is expressed in the asexual blood stages, it is not required for normal asexual development.

**Figure 2.**
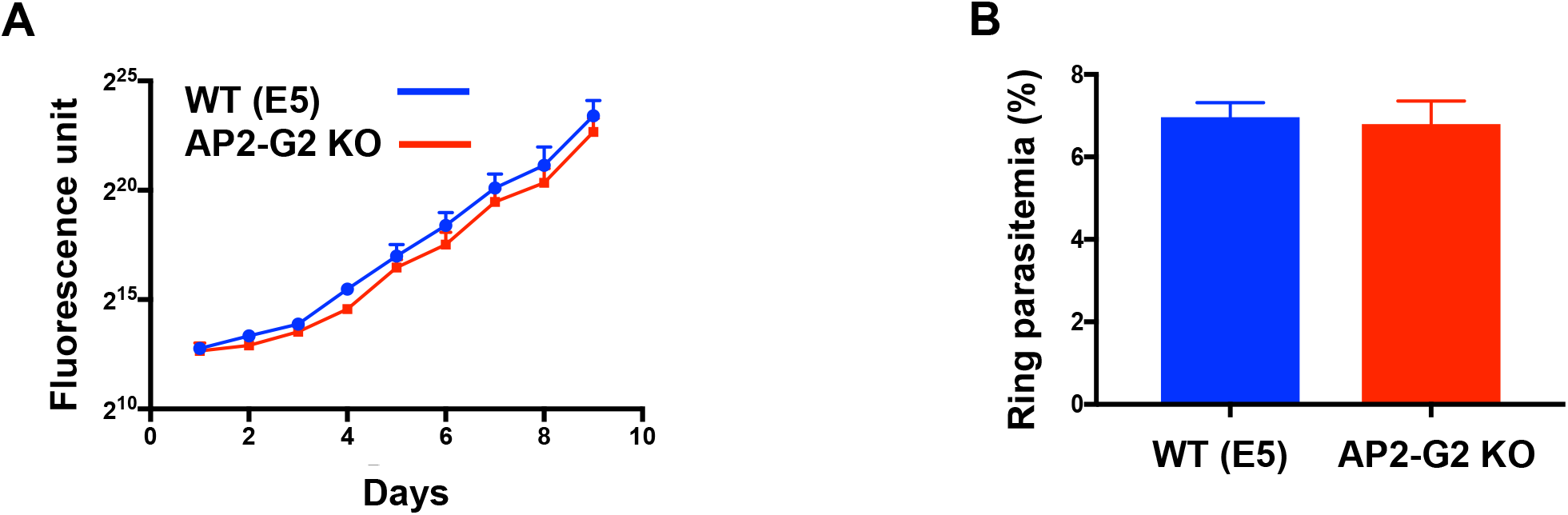
Absence of PfAP2-G2 has no effect on the growth of asexual stage parasite. (A) Growth profiles of the WT and PfAP2-G2 KO parasites measured using an SYBR green assay every 24 hours over a period of 10 days starting with highly synchronous ring-stage parasites. The graph represents the plotted average +/-S.D. of biological triplicates. Fluorescence units are plotted along the Y-axis and the number of days was plotted along the X-axis for the experiment done in triplicate. (B) Bar graph representing the measurement of the multiplication rate of the WT and PfAP2-G2 KO parasites. The values are the average of three replicates, and the error bars represent the standard deviation.

### PfAP2-G2 is expressed in the gametocyte stages

Recent transcriptomic data has shown that the mRNA abundance of *pfap2-g2* in gametocytes is low relative to the asexual stages, with a broad peak seen only in the early gametocyte stages followed by low expression during the later stages (**Supplementary Fig. 3**) (Biljon et al. 2019). To characterize protein expression of PfAP2-G2 during gametocyte development we imaged PfAP2-G2::GFP parasites (see methods). PfAP2-G2 was expressed in all stages of gametocyte development, but it was not confined to the nucleus in later stages (**Figure 3A**). Indeed nuclear fractionation of stage III gametocytes detected expression of full-length protein in both fractions, unlike in asexual parasites (**Figure 3B**). It also seems that the nuclear protein undergoes different proteolytical processing in gametocytes versus the asexual stages.

**Figure 3.**
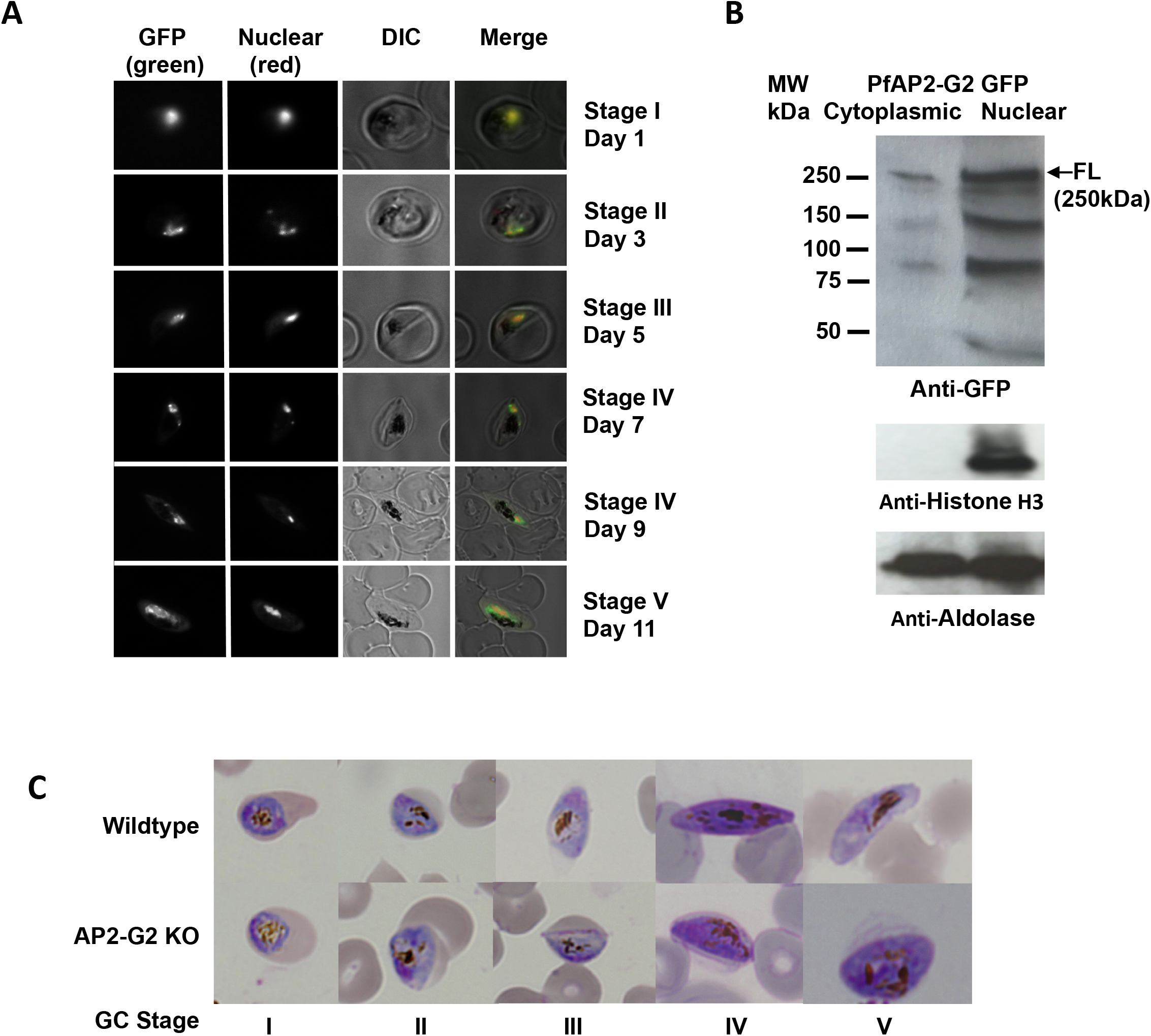
PfAP2-G2 is expressed throughout the gametocyte stages and is essential for maturation of gametocytes. (A) Images obtained by live fluorescence microscopy performed on Stage I through Stage V gametocytes. The DIC image was merged with the DRAQ5 nuclear stain imaging to confirm the nuclear localization of the signal. The results indicate that PfAP2-G2 was expressed in all stages of the gametocytes and was localized to the nucleus. (B) Nuclear (N) and cytoplasmic (C) fractions from the stage III gametocytes were subjected to western blot analysis using anti-GFP antibodies (1:1000). The GFP-tagged AP2-G2 of expected size (~250 kDa) was detected only in the nuclear fraction of the parasite lysate. The WT parasite was used as a negative control. Anti-histone H3 (1:3000) and anti-aldolase (1:1000) were used as the loading controls for the nuclear and cytosolic fractions, respectively. (C) Giemsa-stained smears of gametocytes from the WT and PfAP2-G2 KO lines showed that the morphology of the KO gametocytes appeared normal until Stage III. The KO gametocytes were unable to develop beyond Stage III and showed severe morphological defects.

### PfAP2-G2 is essential for gametocyte maturation

To determine if sexually committed PfAP2-G2 gametocytes mature fully to Stage V, we induced PfAP2-G2 KO and WT parasite lines to undergo gametocytogenesis and followed their development for 14 days, monitoring their maturation via morphological analysis of Giemsa-stained thin-blood smears. Whereas WT E5 parasites matured into Stage V gametocytes, most *pfap2-g2* KO parasites committed to sexual development could not progress beyond Stage III (**Figure 3C**). PfAP2-G2 is thus a critical regulator of gametocyte maturation and development.

### PfAP2-G2 makes extensive genome-wide interactions

Because PfAP2-G2 is predicted to be a DNA-binding protein (Campbell, de Silva, Olszewski, Elemento, & Llinás, 2010; Yuda et al., 2015) we sought to determine the genome-wide localization of PfAP2-G2 in both asexual and gametocyte lifecycle stages. To do this we first performed ChIP-seq at the trophozoite stage using the PfAP2-G2::GFP parasites. Peak calling identified ~5000 peaks in at least 2 of the 3 biological replicates (FDR 0.05) (**Supplementary Fig. 4A**). All replicates showed a high correlation with each other with respect to the overall peaks identified (**Figure 4A**). To our surprise only 401 of the predicted binding sites, corresponding to 120 genes, were found in the non-coding upstream regions of genes (**Supplementary Table 1**), while 4,600 peaks, corresponding to 2,932 genes, were located in exonic gene bodies (**Supplementary Table 2**). This differs from what has been previously observed for other characterized *P. falciparum* ApiAP2 proteins, such as SIP2 (Flueck et al. 2010), AP2-I (Santos et al. 2017) and AP2-G (Josling et al. 2020) which were found to predominantly bind to non-coding upstream regions. Although widescale binding to gene bodies was not previously reported for PbAP2-G2 (Yuda et al. 2015), a re-analysis of the ChIP-seq data from this study using the same pipeline that we used for our ChIP-seq data analysis, revealed that PbAP2-G2 is also broadly distributed across gene bodies in the *P. berghei* genome with 4,345 binding sites.

**Figure 4.**
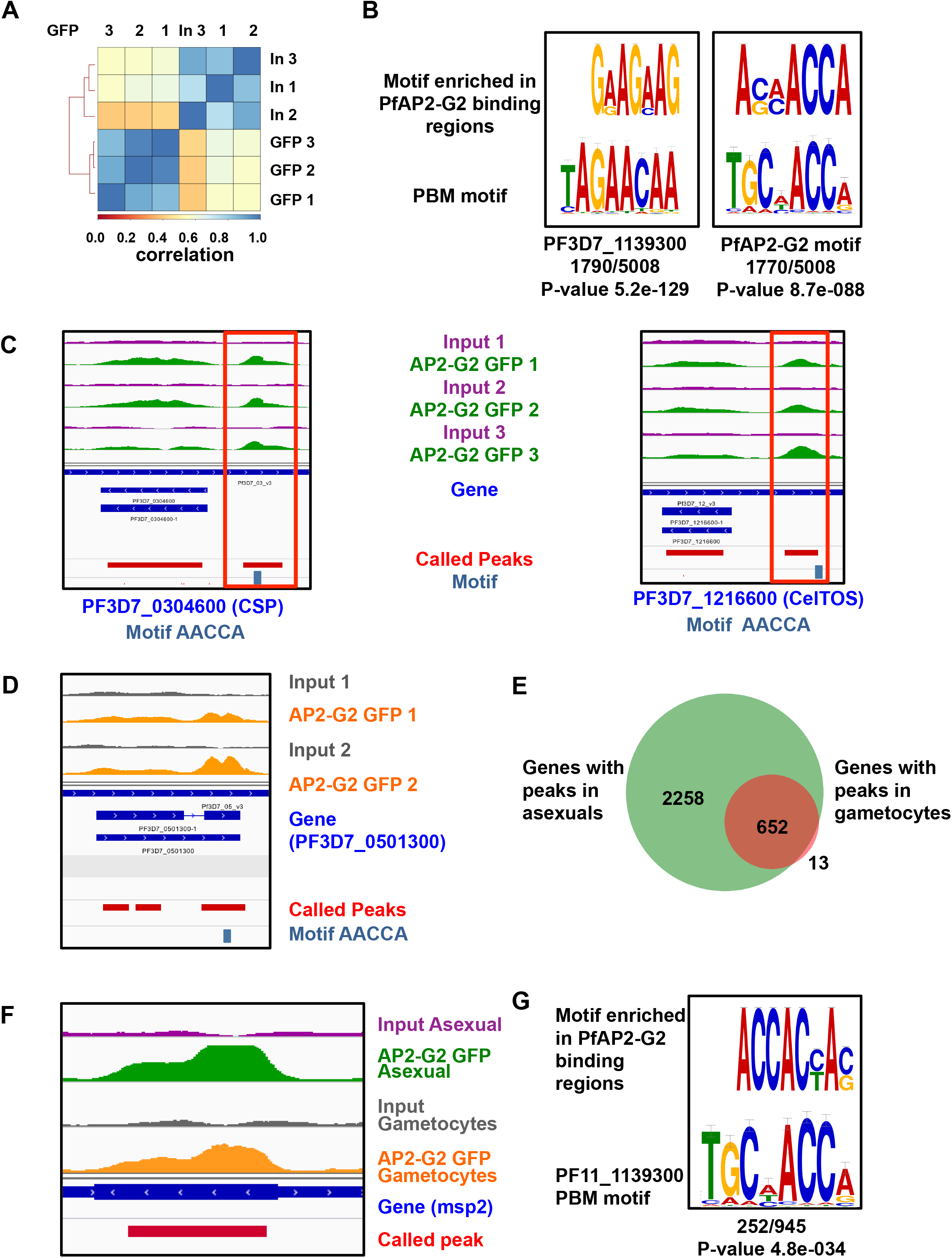
Genome wide predictions of PfAP2-G2 targets and binding regions by ChIP seq. (A) Pearson correlation analysis on three biological replicates of ChIP-seq including the Input (In) and AP2-G2 GFP (GFP) (B) DREME logo for the motif enriched in the genomic locations bound by PfAP2-G2 in the in the gene body and non-coding regions in asexual stage. Shown is the enriched PF11_1139300 motif AGAA and PfAP2-G2 motif ACCA obtained by DREME along with the PBM motif. The P-values are reported by DREME. (C) Example PfAP2-G2 peaks in asexual stages for all three replicates. Peaks are represented as red tracks below with the motif present is highlighted in blue. (D) Example PfAP2-G2 peaks in gametocytes for both replicates. (E) Venn diagram showing common genes with PfAP2-G2 peaks in asexuals and gametocytes (F) Representative peaks for the gene *msp2* enriched with PfAP2-G2 in both asexual and gametocyte stages. (G) DREME logo for the motifs enriched in the genomic locations bound by PfAP2-G2 in the in the gene body and non-coding regions in gametocytes. The P-values are reported by DREME.

An analysis of all the regions (both upstream regions and ORFs) bound by PfAP2-G2, using DREME (Bailey 2011) identified two significantly enriched DNA sequence motifs. The first is an AGAA sequence motif which is related to a previously reported DNA motif found to associate with one (of the three) AP2 domains of the ApiAP2 protein PF3D7_1139300 (Campbell et al. 2010). The second is an ACCA core motif (**Figure 4B**) closely resembling the previously identified PfAP2-G2 binding PBM motif. (Campbell et al. 2010), demonstrating the success of ChIP experiments.

In order to characterize possible regulatory targets of PfAP2-G2, we first focused on the 120 genes that contain PfAP2-G2 peaks in their upstream regions. Nearly 87% of these 120 putative target genes are bound within 2.5kb upstream of the start codon (**Supplementary Fig. 4B**). 278 of the remaining intergenic peaks were localized to the subtelomeric ends of each chromosome (**Supplementary Fig. 4C**). For further analysis we only considered genes with upstream binding interactions less than 2.5 kb from the start codon, leaving 82 candidate genes **(Supplementary Table 1)**. Interestingly, PfAP2-G2 at the asexual trophozoite stage binds to the promoters of genes known to be expressed later in the gametocyte and mosquito lifecycle stages. These include 6-cysteine protein (*p36* PF3D7_0404400), cysteine repeat modular protein 2 (*crmp2* PF3D7_0718300), sporozoite protein essential for cell traversal (*spect* PF3D7_1342500), circumsporozoite protein (*csp* PF3D7_0304600), cell traversal protein for ookinetes and sporozoites (*celtos* PF3D7_1216600), secreted protein altered thrombospondin repeat protein (*spatr* PF3D7_0212600), stearoyl-CoA desaturase (*scd* PF3D7_0511200), and *cap380* oocyst capsule protein (PF3D7_0320400) among others (**Supplementary Table 1**) (Chattopadhyay et al. 2003; van Dijk et al. 2010; Espinosa et al. 2017; Gratraud et al. 2009; Ishino et al. 2004; Itsara et al. 2018; Singh et al. 2007; Thompson et al. 2007; Zhao et al. 2016). Representative PfAP2-G2 peaks upstream of two target genes are shown in (**Figure 4C**). We also found PfAP2-G2 associated with the upstream regions of genes that are highly transcribed in the asexual ring stage (as analyzed in (López-Barragán et al. 2011)) including the knob-associated histidine rich protein (*kahrp* PF3D7_0202000), *Plasmodium* helical interspersed subtelomeric proteins (*phista* PF3D7_0115100), several variant family genes including *vars and rifins*, and genes encoding proteins involved in egress and invasion such as the Rh5 interacting protein (*ripr* PF3D7_0323400) and serine repeat antigen 7 (*sera7* PF3D7_0207400) (Miller et al. 2002; Pei et al. 2005; Sargeant et al. 2006; Volz et al. 2016). Therefore, PfAP2-G2 binds to the promoters and ORFs of genes that are expressed at a different developmental stage than that in which AP2-G2 is expressed, suggesting that, like in the rodent malaria species, this is a transcriptional repressor protein.

We next examined the genome-wide occupancy of PfAP2-G2 in stage III gametocytes by ChIP-seq. We identified 947 peaks in common across both replicates, corresponding to 674 genes at an FDR of 0.05 (**Figure 4D**) (**Supplementary Table 3, Supplementary Table 4)**. PfAP2-G2 binding, once again, was found in both upstream regions and within gene bodies. Although PfAP2-G2 bound many fewer regions in gametocytes compared to asexual trophozoites, which may be associated with the difficulty in obtaining high amounts of chromatin in gametocytes, virtually all stage III PfAP2-G2 target genes were also targets during the trophozoite stages (**Figure 4E**). This suggests that there are a large number of genes that are always bound by PfAP2-G2 during both asexual and sexual parasite development (**Figure 4F**). DNA motif analysis yet again identified the ACCA (**Figure 4G**).

### Genetic disruption of PfAP2-G2 leads to aberrant gene expression in both asexual and sexual life cycle stages

Using synchronized parasites, we collected seven total RNA samples from WT or KO parasites throughout the 48-hour asexual life cycle for transcriptome analysis. In a separate experiment, total RNA samples were collected during sexual differentiation every 12 hours for 7 continuous days starting at 2 days post-gametocyte induction (**Supplementary Fig. 5**). Gene expression analysis of the asexual stage timecourse using Significance Analysis of Microarray (SAM) (Tusher, Tibshirani, and Chu 2001) identified 327 differentially expressed transcripts (>1.5 log2FC, 0.30 FDR) in the PfAP2-G2 KO line. Out of these, 237 showed enhanced transcript abundance in the PfAP2G2 KO line and 90 showed decreased abundance (**Supplementary Fig. 6, Supplementary Table 5)**. This result is quite surprising given the lack of any asexual growth phenotype between the PfAP2-G2 KO and the WT (**Figure 2**), implying that these changes in gene expression do not impact asexual parasite fitness or gametocyte commitment. A comparison of transcript abundance in the KO and WT sexual stages identified increased abundance for 274 transcripts and reduced abundance for 169 transcripts in the PfAP2-G2 KO line (>1.5 log2FC, 0.30 FDR) (**Supplementary Fig. 6, Supplementary Table 6**). These differential transcription patterns presumably lead to the stall in development at Stage III gametocytes in parasites lacking PfAP2-G2.

To determine at which stage the significantly changing genes (by SAM) are maximally transcribed in wild-type parasites, we used a published RNA-seq dataset covering four different asexual stages (rings, early trophozoite, late trophozoite, and schizonts), two gametocyte stages (stage II and stage V), and the ookinete stage using the 3D7 *P. falciparum* strain (López-Barragán et al. 2011). Using this dataset, we identified the 3D7 parasite lifecycle stage at which each gene has the highest reads per kilobase of transcript, per million mapped sequencing reads (RPKM). For the asexual transcriptome, most of the differentially expressed genes we measured as differentially regulated in the PfAP2-G2 KO are normally transcribed maximally in stage V gametocyte (and not in asexual stages), followed by the ookinete and ring stages (**Figure 5A**). This suggests that loss of PfAP2-G2 impacts the expression of genes from late-stage gametocytes, ookinetes, and asexual stages. However the number of genes showing enhanced transcript abundance is strikingly high for genes that are normally expressed in stage V gametocytes (**Figure 5A**). Therefore, PfAP2-G2 may prevent the premature expression of late-stage gametocyte genes during asexual development.

**Figure 5.**
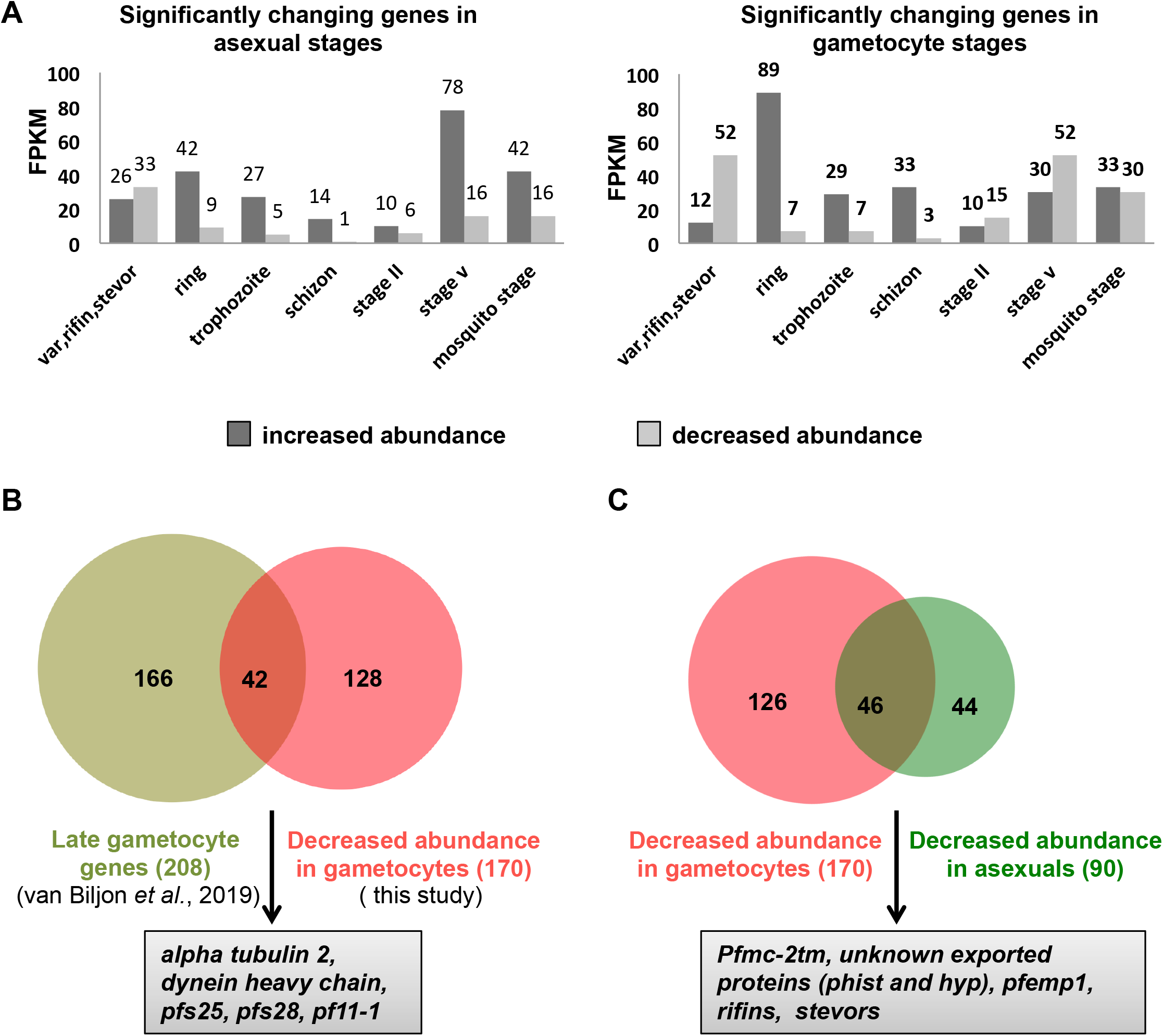
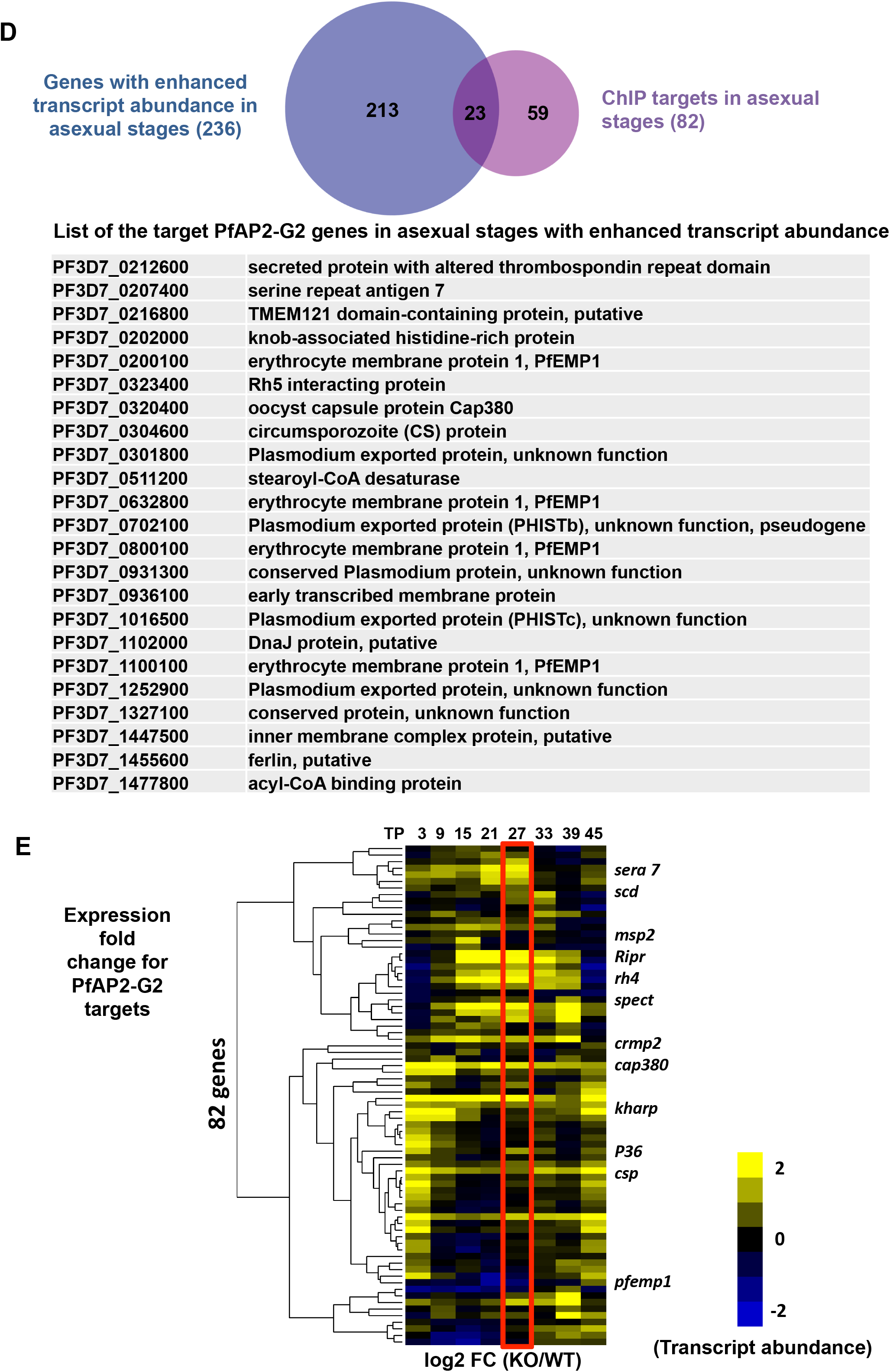
Transcriptional changes in absence of PfAP2-G2. (A) Classification of significantly changing genes in asexual and gametocyte stages based on its maximum transcription in either asexual, gametocytes or mosquito stages, using RNA-seq data (López-Barragán et al., 2011). (B) Overlap between the genes with decreased transcript abundance (>1.5 log2FC, 0.30 FDR) in gametocytes and earlier established late gametocyte markers (van Biljon et al., 2019). (C) Venn diagram showing overlap between genes with decreased transcript abundance in gametocytes and asexual stages. (D) Venn diagram showing number of overlapping genes with PfAP2-G2 enrichment in the upstream regions and also increased transcript abundance (analyzed by SAM, >1.5 log2FC, 0.30 FDR) in absence of PfAP2-G2. (E) Expression fold change (log2 KO versus WT) for the genes that are bound by PfAP2-G2 in the upstream regions in asexual stages. The genes are clustered based on Pearson’s correlation. Boxed is the timepoint at which ChIP-seq was performed.

In the gametocyte transcriptome, we observed the opposite trend when comparing to the López-Barragán data, where 70% of the transcripts that showed increased abundance in the KO line were usually maximally abundant in the asexual stages, especially at the ring stage (**Figure 5A**). Examples of genes whose mRNA abundance was higher in gametocytes include knob-associated histidine-rich protein (*kahrp* PF3D7_0202000), 10 different *Plasmodium* helical interspersed subtelomeric proteins (*phista, phistb, phistc),* 7 different serine/threonine protein kinase *(FIKK family)* proteins, 12genes encoding Plasmodium exported proteins *(hyp2, hyp8, hyp9, hyp10, hyp11, hyp12, hyp16)* merozoite surface proteins (*msp1, msp2, msp6, msp7, msp11*), the ApiAP2 transcription factor *ap2-l* (PF3D7_0730300), serine repeat antigen (*sera5*, PF3D7_0207600) among others (**Supplementary Table 6**) (Pei et al., 2005; Sargeant et al., 2006; Nunes, Goldring, Doerig, & Scherf, 2007; Beeson et al., 2016). These results indicate that PfAP2-G2 represses a subset of genes throughout both asexual and sexual development, and the absence of PfAP2-G2 leads to aberrant timing of gene expression.

To identify enriched DNA sequence motifs associated with gene expression changes, we analyzed the 5’ upstream region (1000bp upstream of the start codon) of the genes showing significant change in transcript abundance in the asexual and sexual stages using the Finding Informative Regulatory Elements (FIRE) algorithm (Elemento, Slonim, and Tavazoie 2007). The most significant enriched sequence motif was the known DNA-interacting motif for PfAP2-G2, TGCAACCA (p-value 1.24 e-21) in genes with enhanced transcript abundance in the asexual stage (**Supplementary Fig. 6**). Interestingly, we also found an AGAACAA (p-value 3.3574e-15) DNA-binding motif that is recognized by the ApiAP2 protein, PF3D7_1139300 (**Supplementary Fig. 6**). Similar motifs were found for genes with increased transcript abundance in gametocytes (**Supplementary Fig. 6**). Intriguingly both *pfap2-g2* and *pf3d7_1139300* (*pf11_0404*) are expressed at similar times throughout asexual development based on transcriptomic data, suggesting that the two proteins may be acting together in some manner to regulate transcription (**Supplementary Fig. 7**). Based on earlier evidence, genetic disruption of the *pf3d7_1139300* orthologue in *P. berghei* (Gene ID) and *P. yoelii* (Gene ID) was refractory indicating its essentiality (Modrzynska et al., 2017; Zhang et al., 2017). Single-cell RNA-seq has shown a sharp upregulation of the gene encoding the ApiAP2 protein PF3D7_1139300 when *ap2-g* expression peaks just before egress in committed schizonts (Poran et al. 2017). Although we couldn’t detect the ACCA PfAP2-G2 binding motif associated with genes showing decreased transcript abundance (**Supplementary Fig. 8**), we found enrichment of the AGACA motif, which has been associated with gametocyte commitment and development (**Supplementary Fig. 8**) (Young et al., 2005; Bischoff & Vaquero, 2010; Painter, Carrasquilla, & Llinás, 2017), in genes showing decreased transcript abundance in gametocytes.

Genes with decreased mRNA abundance in the gametocyte transcriptome from the PfAP2-G2 KO line (170 genes) overlap significantly with genes previously reported as late gametocyte markers (Van Biljon et al. 2019) (cluster 9, 208 genes) (**Figure 5B**). Some example genes are alpha tubulin 2 (PF3D7_0422300), TRAP-like protein (*tlp* PF3D7_0616500), secreted ookinete protein (*psop13* PF3D7_0518800, *psop20* PF3D7_0715400), dynein heavy chain (PF3D7_0729900), ookinete surface protein P28 (PF3D7_1031000), ookinete surface protein P25 (PF3D7_1031000) (Rawlings et al., 1992; Heiss et al., 2008); Ecker, Bushell, Tewari, & Sinden, 2008; Villard et al., 2007; Duffy & Kaslow, 1997). Therefore, one reason that PfAP2-G2 KO parasites may fail to progress beyond stage III may be due to the aberrant expression of mRNAs required during this and subsequent stages of gametocyte development. The inhibition of gametocyte maturation may also be due to the decreased expression of a few critical genes required for gametocyte development, such as male development gene 1 (*mdv1* PF3D7_1216500), gametocyte erythrocyte cytosolic protein (*geco*, PF3D7_1253000), gametocyte exported protein (*gexp06* PF3D7_0114000)*, and* gametocyte specific protein (*pf11-1* PF3D7_1038400) during the asexual stage. Interestingly, we also see overlap between the genes with decreased transcript abundance in both the asexual and gametocyte stages. These overlapping genes are enriched mostly for members of the *var*, *stevor*, and *rifin* variant gene families, *Pfmc-2tm* among others (**Figure 5C**). An early gametocyte proteome has revealed that exported and erythrocyte remodeling proteins are the most overrepresented proteins in gametocytes (Silvestrini et al. 2010). Although members of the PfEMP1, STEVOR and RIFIN protein repertoire are expressed in gametocyte stages, their role is not well established (Sharp et al., 2006; Petter, Bonow, & Klinkert, 2008; Tibúrcio et al., 2012, Mwakalinga et al., 2012; Tibúrcio et al., 2013; Neveu et al., 2018; Neveu & Lavazec, 2019;), although decreased abundance of these genes may contribute to the phenotypic effect.

### PfAP2-G2 binding in the promoter and in gene bodies is associated with differential regulation of transcription

Out of 82 genes with upstream PfAP2-G2 peaks, only 23 displayed significantly elevated mRNA abundance (as analyzed by SAM, >1.5 log2FC, 0.30 FDR) (**Figure 5D**). This suggests that the transcription of other genes may be impacted indirectly in the PfAP2-G2 knockout. On the other hand, none of the genes with significantly reduced mRNA abundance were bound by PfAP2-G2 in the upstream regions. However, considering the stage at which ChIP was performed we see that the mRNA abundance of many of the PfAP2-G2 ChIP-seq targets do not change in the absence of PfAP2-G2 indicating that binding of PfAP2-G2 alone might not be sufficient to affect transcription (**Figure 5E**). PfAP2-G2 binding to gene bodies results in both repression and activation in the *pfap2-g2* KO. Open reading frames bound by PfAP2-G2 in the gene-body in gametocytes showed similar effects (**Supplementary Fig. 9**). Overall these results indicate that PfAP2-G2 binding to the upstream promoter region is largely associated with repression, but PfAP2-G2 alone is not sufficient to repress transcription.

### PfAP2-G2 shares occupancy with exonic epigenetic marks during asexual development

To understand the role of the pervasive genome-wide PfAP2-G2 binding in exonic gene bodies detected by ChIP-seq (**Supplementary Table 2**), we investigated whether PfAP2-G2 is associated with any known epigenetic marks. We calculated a pairwise Pearson correlation coefficient by comparing the read counts per 1000bp bins of the PfAP2-G2 ChIP-seq results against an array of histone marks for which ChIP data is available (H3K9me3, H4ac, H4K20me3, H3K26ac, H3K9ac, H3K4me3 (Karmodiya et al., 2015), H3K36me2 and H3K36me3 (Jiang et al., 2013), as well as heterochromatin protein 1 (PfHP1) (Brancucci et al., 2014)). Interestingly, we found that PfAP2-G2 occupancy has the strongest correlation with H3K36me3 (R=0.933), followed by H4K20me3 (R=0.911), H3K27ac (R=0.826), and H4ac (R=0.765) (**Figure 6A**), which are all histone marks that show moderate positive correlation with *P. falciparum* transcription and are associated with transcriptionally poised genes (Karmodiya et al. 2015). In total, 3,633 (80%) PfAP2-G2 binding sites co-occur with H3K36me3 marks (**Figure 6B, 6C**). The functional role of the association between PfAP2-G2 and H3K36me3 is, and what the potential role of PfAP2-G2 in H3K36me3 recruitment remains to be determined. PfAP2-G2 also co-occurs with PfHP1 towards the end of chromosomes although this correlation is moderate (R=0.6) (**Figure 6D, 6E**).

**Figure 6.**
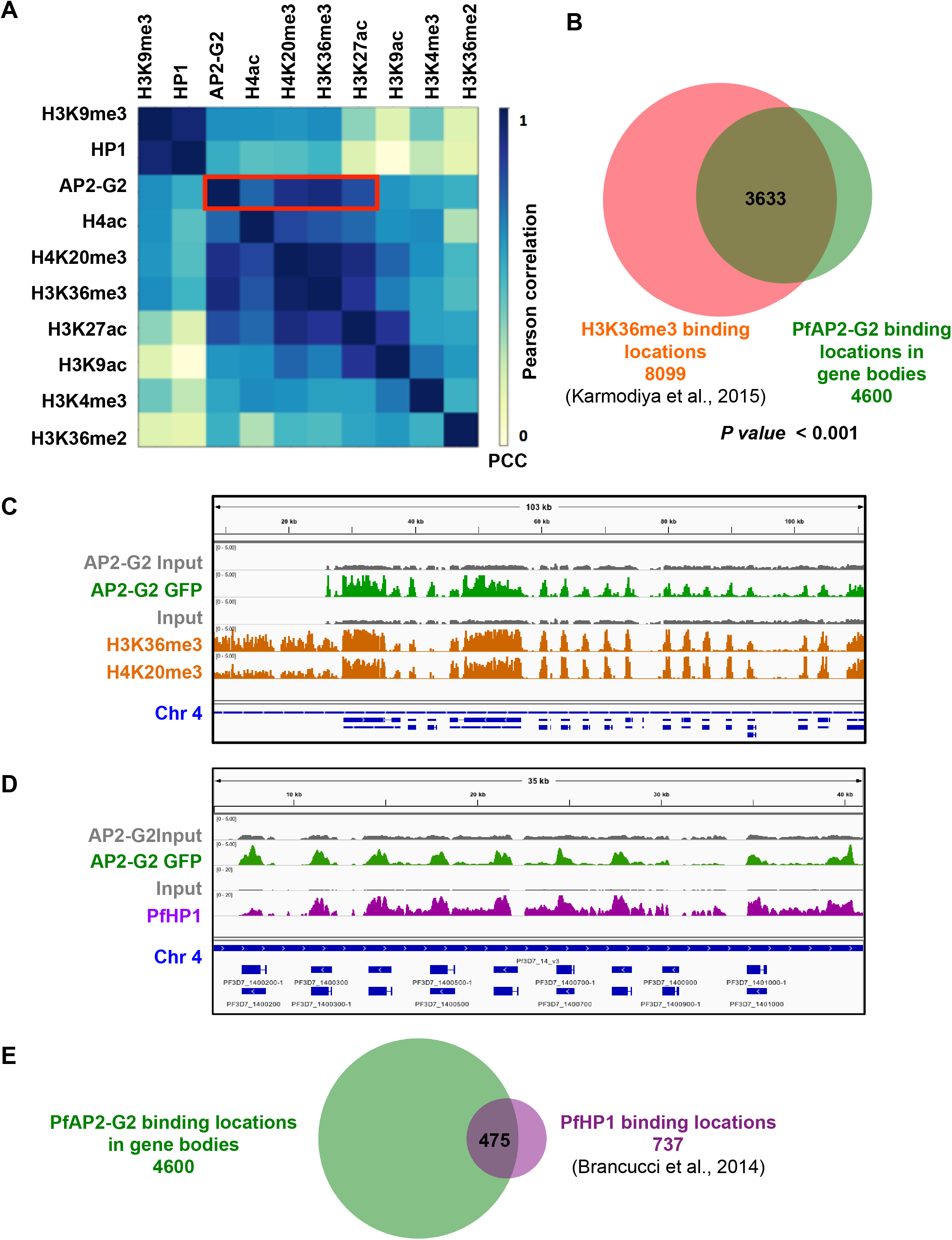
PfAP2-G2 shows high correlation with H3K36me3 repressive histone modifications. (A) Heatmap showing correlation between occupancy of PfAP2-G2 and nine histone modifications in the gene body of all P. falciparum genes (n=5735). PfAP2-G2 strongly correlates with H3K36me3 and H4K20me3 (B) Venn diagram comparing the peaks obtained by ChIP-seq on H3K36me3 marks (oange) (Jiang *et al.*, 2013) and PfAP2-G2-GFP (green) shows that 80% of times PfAP2-G2 shows co-occupancy with H3K36me3 marks. (C) IGV screenshot showing co-occupancy of PfAP2-G2 with H3K36me3 and H4K20me3. First gray track represents Input for PfAP2-G2 and second green track shows the regions bound by PfAP2-G2. Third gray track represents input for H3K36me3 and H4K20me3 and the last two orange tracks represent the regions bound by H3K36me3 and H4K20me3 as indicated in the figure. Blue track shows the genes. (D) IGV screenshot showing co-occupancy of PfAP2-G2 with PfHP1. First gray track represents Input for PfAP2-G2 and second green track shows the regions bound by PfAP2-G2. Third gray track represents input for PfHP1 and the last two pink tracks represent the regions bound by PfHP1 as indicated in the figure. Blue track shows the genes. (E) Venn diagram comparing the peaks obtained by ChIP-seq on PfHP1 marks (pink) (Brancucci *et al*., 2014) and PfAP2-G2-GFP (green) shows that 80% of times PfAP2-G2 shows co-occupancy with H3K36me3 marks.

### PfAP2-G2 interacts with the chromatin remodeling machinery

To identify proteins interacting with PfAP2-G2, we performed immunoprecipitations (IP) from PfAP2-G2::GFP parasites at the trophozoite stage using an anti-GFP antibody. Western blot analysis confirmed that the full length PfAP2-G2 protein was purified and eluted (**Figure 7A**), although smaller GFP-positive band(s) were also seen. The resulting IP was separated by SDS PAGE and two regions of the gel (upper and lower band, **Figure 7A**) were subjected to protein analysis by mass spectrometry. The purpose of analyzing these bands separately was to ensure that we could detect all of the proteins including the low abundance protein which otherwise would get masked when analyzing all of the bands together (**Figure 7B**).

**Figure 7.**
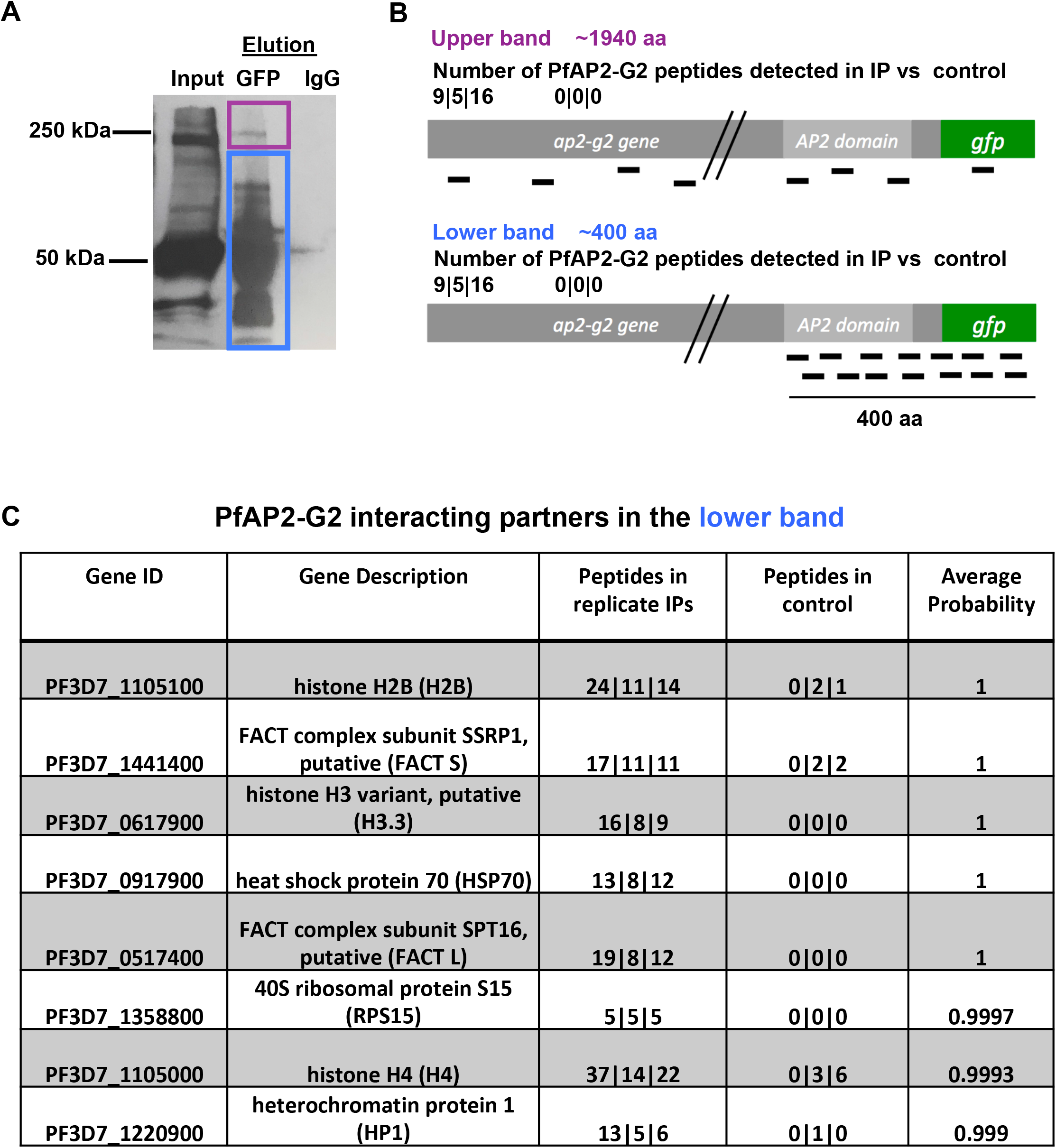
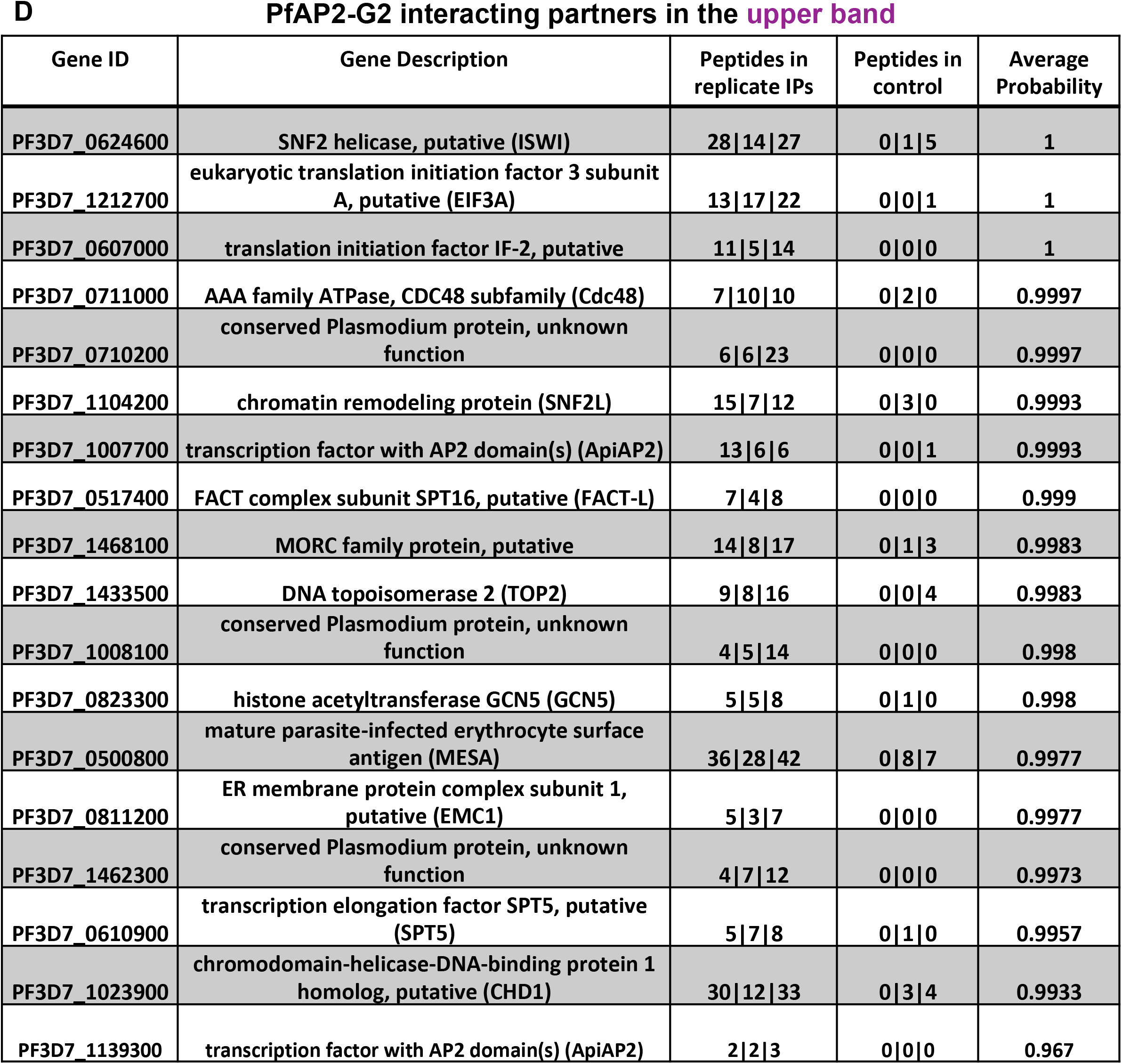
Identification of potential interacting partners of PfAP2-G2. (A) SDS PAGE showing immunoprecipitation (without crosslinking) of the PfAP2-G2 GFP parasite line using antibodies against GFP. Full length PfAP2-G2 and the cleaved product was analyzed separately by mass spectrometry to identify the interacting partners of PfAP2-G2. (B) Schematic showing the peptides recovered from upper and lower band. Also shown are the number of PfAP2-G2 peptides in both the bands and in the controls. (C) List of PfAP2-G2 interacting partners in the lower band with high probability of interaction measured by SAINT (pSAINT 0.9 and 1%FDR). (D) List of PfAP2-G2 interacting partners in the upper band with high probability of interaction measured by SAINT (pSAINT 0.9 and 1%FDR) in the upper band.

The proteomic IP data was analyzed using SAINT (Choi et al. 2011) with a score of 0.9% and 1% FDR across all three replicates (**Supplementary Table 7**). Interestingly, we identified HP1 as an associated protein, which is consistent with the observed co-occurrence of PfAP2-G2 peaks with HP1 (**Figure 6D**) **(Figure 7C)**. The upper band contained larger chromatin remodeling proteins such as ISWI, SNF2L, GCN5, FACT complex proteins (FACT-S and FACT-L), a MORC family protein, and several ApiAP2 family proteins, including PF3D7_1139300 (PF11_0404) whose motif was found to be enriched in both the transcription and ChIP-seq data **(Figure 7D)**. This suggests that direct interaction of PF3D7_1139300 and PfAP2-G2 may be required for the regulation of a subset of target genes.

## Discussion

Gametocytogenesis in *Plasmodium falciparum* parasites is a long 10-12 day developmental progression resulting in the formation of fertile male and female gametes that will form an oocyte only upon transmission to the mosquito host. Recent studies have shed light on the commitment and differentiation of asexual parasites into sexual parasites, which is driven by the AP2-G master regulator (Josling et al. 2020), (Kafsack et al. 2014). However, how asexual cells are reprogrammed for gametocyte commitment, and the downstream post-commitment regulation of sexual development is not well characterized. There are known examples of gametocyte stage genes such as *pfsegxp, mdv1,* and *pfs16* that are expressed very early on in the asexual stages suggesting that programming for gametocytogenesis may begin early in asexual development, and disruption of this program could result in profound effects post-commitment during sexual development (Furuya et al., 2005; Joice et al., 2014; Painter et al., 2017; Nixon et al., 2018).

In this study we show that the ApiAP2 protein PfAP2-G2 is essential for the development and maturation of *P. falciparum* gametocytes. PfAP2-G is expressed from the very early trophozoite stage through to stage V gametocytes. This is in contrast to *P. berghei*, where the protein is expressed from 16hpi, during the schizont stage of development (Yuda et al. 2015). Live fluorescence microscopy and nuclear fractionation show that PfAP2-G2 is localized to the nucleus of the trophozoite, schizont, and early gametocyte stages (**Figure 1B**). In later stage gametocytes PfAP2-G2 protein levels are reduced and expressed throughout the cytoplasm (**Figure 3A**). This is consistent with the mRNA abundance profile, demonstrating peak *pfap2-g2* expression in the early asexual stages, whereas in gametocytes its expression is broad and weak (Biljon et al. 2019). The nuclear localization and strength of protein expression indicate that the major role for PfAP2-G2 is during the asexual stages and in early gametocytes.

By performing genome-wide ChIP-seq analysis at the asexual trophozoite stage and stage III gametocytes, we show that PfAP2-G2 extensively binds to both intergenic and genic regions. Genes bound by PfAP2-G2 in the upstream regions are largely expressed later in the gametocyte and mosquito stages. However, PfAP2-G2 also binds the promoters of genes expressed in the asexual ring stage and genes that are transcribed in schizont stages like *msps* encoding the merozoite surface proteins, which are regulated by PfAP2-I (Santos et al. 2017). Of course MSPs are unlikely to be required in committed gametocytes, as they no longer egress from the red blood cell for subsequent re-invasion. Interestingly, genes bound by PfAP2-G2 in stage III gametocytes are essentially the same as those bound in trophozoites. An analysis of the DNA motifs enriched in the regions bound by PfAP2-G2 identified two motifs: an AGAA motif and an ACCA motif (**Figure 4B** and **5B**). The latter resembles the PBM-based predicted motif for PfAP2-G2, but it is different from that identified in *P. berghei* (GTTGT), even though the AP2 DNA-binding domain is highly conserved (97% identical). We also found enrichment for a second motif in the PfAP2-G2-bound regions, which resembles the *in vitro* DNA sequence motif bound by the ApiAP2 protein PF3D7_1139300 (PF11_0404) (Campbell et al. 2010). Supporting the hypothesis that these ApiAP2 proteins may function together in a complex, our immunoprecipitation experiments against PfAP2-G2 GFP detected AP2 PF3D7_1139300 (PF11_0404) as an interaction partner (**Figure 7**, **Supplementary Table 7**).

Parasites expressing a truncated PfAP2-G2 version proliferate normally in the asexual stages. These results are surprising given the many global transcriptional (327 transcripts, >1.5 log2FC, 0.30 FDR) changes occurring in the asexual stages, and suggests that the parasite is tolerant to large fluctuations in gene expression. Compared with wild-type parasites, the expression levels of 90 genes (>1.5 log2FC, 0.30 FDR) were significantly decreased during asexual development in PfAP2-G2 KO parasites, including genes that have been reported to have functional roles in gametocytes, perhaps leading to the developmental stall observed in Stage III gametocytes (Painter, Carrasquilla, & Llinás, 2017; Van Biljon et al., 2019). This suggests that the program for sexual progression is established very early, in the asexual stage. KO of PbAP2-G2 also resulted in the downregulation of gametocyte genes (Yuda et al. 2015). Genes upregulated in the PfAP2-G2 KO were mostly those normally expressed at other later developmental stages including the gametocyte and ookinete stages (**Figure 5A**). In the upstream promoter regions of genes showing increased transcript abundance in the PfAP2-G2 KO, we found an enrichment of the PfAP2-G2 ACCA motif. PfAP2-G2 therefore is involved in the repression of a number of genes that are not required for asexual development.

A previous study that examined the genome-wide binding of AP2-G2 from *P. berghei* gametocytes using ChIP-seq reported PbAP2-G2 binding at over 1,500 genes that were mostly associated with roles in asexual-stage proliferation (Yuda et al. 2015). Transcriptome analysis in gametocytes revealed the upregulation of 927 genes by more than twofold in *pbap2-g2* KO lines, suggesting that PbAP2-G2 acts as a repressor for asexual-stage genes during sexual development. However, only 397 (26.5%) of the PbAP2-G2 bound genes were upregulated in the *pbap2-g2* KO, leading Yuda *et al.* to hypothesize that relieving repression was not sufficient for the upregulation of the rest of the genes. Also, in our data, not all genes that are bound by PfAP2-G2 show enhanced transcript abundance (only 23/82 genes with upstream PfAP2-G2 peaks (28%)) displayed significantly elevated mRNA abundance. In fact, the PfAP2-G2 motif is only present in the promoter of genes upregulated in both developmental stages indicating that, as in *P. berghei*, PfAP2-G2 occupancy is not always predictive of changes in gene expression, perhaps because it requires interaction with other proteins. Once more reflecting the repressive nature of PfAP2-G2, none of the genes bound by PfAP2-G2 in the upstream region are downregulated in the KO.

The effect of PfAP2-G2 truncation in transcript abundance is more pronounced in gametocytes. Unfortunately, due to the massive dysregulation of mRNA transcripts, it is difficult to ascertain which genes cause the stall in development observed at Stage III gametocytes. The significantly upregulated genes in gametocytes are mostly thought to be important for asexual stage development, similar to what was shown in *P. berghei* (Yuda et al. 2015). Interestingly, the ApiAP2 protein AP2-L, which plays a critical role in liver-stage development, is upregulated in the gametocyte stage in both the *P. falciparum* and *P. berghei ap2-g2* knockouts, and parasites lacking PbAP2-G2 are unable to cause liver infection.

We observed a moderate to strong correlation between PfAP2-G2 occupancy and that of H3K9me3 (R=0.933), followed by H4K20me3 (R=0.911), H3K27ac (R=0.826), and H4ac (R=0.765) as well as PfHP1 (R=0.6) (**Figure 6A**). H3K36 methylation is a well-studied mark in model organisms, and carries out several roles. Studies in metazoans have shown that H3K36me3 is enriched at gene exons and plays a role in the regulation of alternative splicing (Kolasinska-Zwierz et al. 2009). Although H3K36me3 correlates with active transcription, another study showed the association of this modification with facultative heterochromatin (Chantalat et al. 2011). Thus, H3K36me3 is involved with both actively transcribed as well as silenced regions and may contribute to the formation of heterochromatin in combination with other histone modifications. At a subset of heterochromatic genes, binding to HP1a is required for enzyme-mediated H3K36me3 demethylation suggesting that there are different ways in which H3K36me3 regulates chromatin (Lin et al. 2012). In *P. falciparum*, H3K36me3 is present along the entire gene body of silent *var* genes including the transcription start site (TSS), and it is deposited by the *P. falciparum* variant-silencing methyltransferase containing a SET domain (PfSETvs) (Jiang et al. 2013). However, in that study enrichment of H3K36me3 was found at the 3’ end of other ring-stage active genes besides the variable antigen *var, rifin* and *stevor* genes in PfSET KO parasites. This result indicates there is an alternative methyltransferase resulting in H3K36 methylation. We speculate that PfAP2-G2 plays a role in regulating regions of the genome which need to be silenced to prevent spurious transcription.

We also found that PfHP1 localized with PfAP2-G2 towards the end of the chromosomes and is associated with both sub-telomeric and chromosome-internal *var* genes and Plasmodium-exported proteins. A direct interaction between PfHP1 and PfAP2-G2 was supported by IP-MS (**Figure 7C**). Another association of PfAP2-G2 that was significant is with the MORC protein (**Figure 7D**) which has been identified to play important roles in chromatin compaction in plants and animals and was recently established as the upstream transcriptional repressor of sexual commitment in *Toxoplasma gondii* (Farhat et al. 2020). This study found that MORC forms a complex with *T. gondii* ApiAP2 transcription factors at sexual stage genes. Two of the *T. gondii* ApiAP2 proteins found to associate with MORC by IP-MS are TgAP2IV-2 and TgAP2IX-9, whose AP2 DNA-binding domains are most identical to the PfAP2-G2 AP2 domain (Farhat et al. 2020). We also found the *P. falciparum* ISWI chromatin-remodeling protein associated with PfAP2-G2, which was recently identified in a complex with the MORC protein at *var* gene promoters (Bryant et al. 2020)(**Figure 7D**).

Previous studies have shown expression of the master gametocyte regulator *pfap2-g* peaks at two times of asexual development (Poran et al. 2017). The initial expression peak in trophozoites is associated with the expression of genes encoding ISWI, SNF2L expression and the ApiAP2 encoding gene *pf3d7_1222400.* The second increase in abundance occurs just before egress in committed schizonts and leads to the expression of genes encoding LSD2, HDA1 and the AP2-G2 associated ApiAP2 protein PF3D7_1139300 (Poran et al. 2017). We propose that PF3D7_1139300 may act in a complex with PfAP2-G2 to compress chromatin structure. In summary, we propose that PfAP2-G2 plays an important role in gene repression during the maturation of the *P. falciparum* parasite sexual stages by associating with other chromatin-related proteins.

## Materials & Methods

### Construction of plasmid for tagging 3’end of *pfap2-g2* and *pfap2-g2* gene disruption

The pDC2-cam-CRT-GFP (Fidock 2000) plasmid was used to tag the 3’ end of the gene *pfap2-g2* with *gfp*. A 948 bp homologous fragment from the 3’ end of the gene was amplified from 3D7 genomic DNA using the oligonucleotide pairs (Forward Primer P7 and Reverse Primer P8, see primer list), which also encoded restriction sites BglII and XhoI for cloning. This fragment was ligated into the BglII-and XhoI-digested pDC2-cam-CRT-GFP resulting in the final vector pDC2-cam-AP2-G2-GFP.

To create the *pfap2-g2* disrupted line, the *ap2-g2* gene locus was targeted using the selection linked integration (SLI) method (Birnbaum et al. 2017). To do this a 500 bp fragment of the 5’ end of *pfap2-g2* was amplified using a forward primer with a NotI restriction site as well as a single point mutation in the homologous sequence to introduce a stop codon (Forward Primer P9) and a reverse primer with an MluI site (Reverse Primer P10) to be ligated into the final vector. This fragment was cloned (in frame) into pSLI-TGD using NotI and MluI restriction sites resulting in the final vector, pSLI-TGD-AP2-G2.

### Culturing, synchronization, transfection, and cloning of *Plasmodium falciparum* parasites

*Plasmodium falciparum* parasites were cultured as previously described (Trager & Jensen, 1976) and maintained in 6% O_2_ and 5% CO_2_ and grown in sterile filtered RPMI 1640 media (Gibco) supplemented with 2 M HEPES, NaHCO_3_ (2g/L), 0.1 M hypoxanthine, 0.25%(w/v) Albumax II (Gibco), and gentamycin (50 ug/ml). The parasites were maintained at a hematocrit of 3% and were kept between 2-5% parasitemia or greater based on the needs of the individual experiments. Parasite synchronization was performed using 5% sorbitol (Lambros and Vandenberg 1979).

To generate transgenic AP2-G2::GFP and AP2-G2 KO lines, transfections were carried out according to a standard protocol (Deitsch, Driskill, and Wellems 2001). The plasmid pDC2-cam-AP2-G2-GFP was transfected into the 3D7 strain of *P. falciparum* whereas pSLI-TGD-AP2-G2 was transfected into the 3D7 clone, E5 (Rovira-Graells et al. 2012). In brief, 100 μg of plasmid DNA (maxi prepped using Qiagen kit) was preloaded into O+ RBCs at 50% hematocrit in cytomix prior to culturing with parasite-infected RBCs. After reinvasion, media containing 2.5 nM WR99210 was added to select for transgenic parasites. Stable integrants were maintained under constant drug pressure and were PCR-verified using specific primers (P1, P2, P4, P5, P6) outside the homology region after the isolation of gDNA (DNeasy Blood and Tissue kit, Qiagen). The resulting PfAP2G2::GFP and *pfap2-g2* ko parasites were cloned by limiting dilution (Rosario 2008) and verified by diagnostic PCR (Supplementary Fig. 1 & 2A) and whole genome sequencing. Sequencing analysis was done by creating a reference genome with GFP and rest of the plasmid at the expected location and reads were aligned to it.

### *Plasmodium falciparum* gametocyte production

Gametocytes were generated using the nutrient deprivation induction method as described (Miao et al. 2013). Briefly, synchronized trophozoites were cultured at ~2% parasitemia using 50% fresh RBC at 4% hematocrit. When the parasitemia reached 7–10%, gametocyte production was induced by nutrient deprivation. The following day, stressed schizonts were split 50% into two flasks by adding fresh blood and fresh media. The following day, ring-stage committed gametocytes were counted as Day 1 of gametocytogenesis. To prevent the reinvasion and propagation of asexual parasites, parasites were treated with media with heparin (20 U/ml) from Day 1 post induction through Day 4. The media was changed daily to ensure the proper development and growth of the gametocytes.

### Nuclear and cytoplasmic protein extraction from parasites and western blotting

Nuclear and cytoplasmic extracts from parasites were prepared as previously described (Voss et al. 2002) using 5-7% of the synchronized parasites. The proteins from the nuclear and cytoplasmic fractions were run on a gel (mini PROTEAN precast TGX gels) and then transferred to a nitrocellulose membrane. The membrane was then blocked using 5% non-fat milk powder in 1X PBS-0.05% Tween for 45 minutes at room temperature. The membrane was then incubated overnight at 4°C with an anti-GFP (ROCHE Anti-GFP, from mouse IgG1K, Millipore Sigma) (1:1000), anti-histone (Anti-Histone H3 antibody, AbCam) (1:3000), anti-aldolase (Anti-Plasmodium aldolase antibody (HRP), AbCam) (1:1000) primary antibody in the blocking buffer. After incubation, the membrane was washed three times with 1X PBS-0.05% Tween and was incubated with the appropriate secondary antibodies, then washed. The secondary antibodies used were as follows: peroxidase-conjugated goat anti-rat HRP conjugate (Millipore) / goat anti-rabbit HRP conjugate (Millipore) / goat anti-mouse HRP conjugate (Pierce) was used (1:3000). Bound antibodies were then visualized on a film after enhancing the signal with ECL (Pierce) as the substrate.

### Parasite growth assay

To compare the growth rates of *pfap2-g2* KO and WT parasites we used a SYBR green growth assay (Vossen et al., 2010). Parasites were sampled every 24 hours for 10 days, starting at 0.1% parasitemia at the trophozoite stage. Both the WT and PfAP2-G2 knockout strains were synchronized using 5% sorbitol (described above) 3x before the start of the timecourse. Parasites were seeded in a 25 cm^2^ culture flask at 0.1% parasitemia and 3% hematocrit. To evaluate growth differences every 24 hours for 10 days, 100 μl of parasite culture was collected in triplicate, transferred to a 96-well plate, and stored at −80°C. Following the completion of the timecourse, the parasites were thawed at room temperature, 100 μl of SYBR green I (Molecular Probes, Eugene Oregon) was diluted into parasite lysis buffer (0.2 μl of SYBR Green I/ml of lysis buffer) and added to each well and mixed. The plates were incubated at 37C for 2 hours. Cell growth was measured as quantification of DNA by SYBR green I using a Tecan GENios microplate detection device at the excitation wavelength and emission wavelength of 485 nm and 535 nm, respectively. All values were plotted as an average of three technical replicates (±S.D.) using GraphPad Prism (version 7).

### RNA purification and cDNA synthesis for DNA microarrays

Total RNA was extracted from tightly synchronized PfAP2-G2::KO and WT parasites every 6 hours, starting at 3 hpi, throughout the 48 hour asexual life cycle. A separate timecourse was designed to collect RNA from gametocytes starting at Day 1 and every 12 hours for the following 7 days. RNA was extracted from parasite-infected RBCs collected at each timepoint using TRIzol (ThermoFisher Scientific), following the manufacturer’s protocol. cDNA synthesis and DNA microarray analysis was carried out as previously described (Painter et al. 2013). Two-channel Agilent DNA microarrays (AMADID 037237) were hybridized, washed, and scanned on an Axon 4200A scanner. Signal intensities for each gene were extracted from the scanned image using Agilent Feature Extraction Software version 9.5. Detailed microarray protocols can be found at http://llinaslab.psu.edu/protocols/. The data were analyzed by Significance Analysis of Microarrays(SAM) (Tusher et al. 2001) for the significantly (>1.5 log2FC, 0.30 FDR) changing genes between the WT and AP2-G2 KO lines using two-class paired t-test throughout the time course. All heat maps were clustered using Cluster 4.0 and was visualized using Java Tree View.

### Chromatin immunoprecipitation and Library preparation for sequencing (ChIP-Seq)

ChIP was performed as described (Josling et al. 2020) using synchronized trophozoite and gametocyte stage PfA2-G2-GFP parasites (around 7% parasitemia). Extracted chromatin was sheared in SDS lysis buffer (200 μl 1× 10^9^ trophozoites and 5×10^8^ gametocytes) to obtain a fragment size of 100-150bp using an M220 focused-ultrasonicator (Covaris Inc.) using the following settings: peak power 75W, 2% duty factor, 200 cycles per burst and total treatment time of 600 s. Sonication time was optimized for PfAP2-G2 using the crosslinked DNA and shearing it for 0, 4, 8, 10, 12 and 15 minutes and then running it on agarose gel for the size estimate. For immunoprecipitations, sheared chromatin was pre-cleared using 20 μl/ml of magnetic beads A/G (Millipore 16-663) for 2 hours at 4°C with gentle agitation. The supernatant was collected and 50μl of the aliquot was removed as input. The rest of the chromatin was incubated overnight at 4°C with 1 μg of polyclonal anti-GFP antibody (Abcam ChIP grade, Abcam 290) or, as control, the same amount of IgG. The immunoprecipitated antibody/chromatin complex was collected using Protein A/G magnetic beads (Millipore 16-663). The input and ChIP samples were reverse cross-linked overnight at 45°C using 0.2M final concentration of NaCl. The samples were then treated with RNAase (30min at 37°C) and proteinase K (3 μL of 20 mg/mL with 2 hours incubation at 45°C) and purified by using the QIAGEN MinElute PCR purification kit. Library was prepared as described earlier (Santos et al. 2017). The final library was quantified using a Qubit fluorometer HD DNA kit and analyzed using an Agilent DNA 1000 Bioanalyzer to assess the quality, size distribution, and detection of any artifacts. Sequencing was performed using an Illumina Hiseq 2500 to obtain 150 bp single-end reads.

### ChIP-seq data analysis

ChIP-seq data analysis was done using tools in Galaxy (usegalaxy.org). Before starting the analysis, the quality of the sequencing reads were assessed using FastQC and were trimmed to remove adapter and low quality bases using Trimmomatic (Bolger, Lohse, and Usadel 2014). The resulting reads were mapped to the *P. falciparum* genome (Pf 3D7 v28, obtained from PlasmoDB) using BWA-MEM (Li and Durbin 2009) and duplicates reads were removed using SAMtools (Li et al. 2009). Mapped sequences were converted into bigwig files using bamCoverage (Ramírez et al. 2016) and were viewed in using the Integrative Genomics Viewer (IGV) (Thorvaldsdóttir, Robinson, and Mesirov 2013) for each input and treatment. MACS2 (Feng et al. 2012) was used to call peaks with q-value cutoff of 0.05. The common overlapping intervals between replicates were established using Multiple Intersect function of BEDtools, and the overlapping intervals were combined into a single file using MergeBED (Quinlan and Hall 2010). The genes closest to the peak summits were identified using closestBED. Peaks that were farther than 2.5Kb were removed from the analysis. When the peak was present between two genes then only the gene closest to the peak was considered for the analysis.

Peak sequences were extracted using extract genomic DNA tool using genomic coordinates, and motifs associated with called peaks were identified using DREME (Bailey 2011) and compared to identified ApiAP2 motifs using TOMTOM (Gupta et al. 2007). The correlation heatmap of the replicates performed on GFP tagged PfAP2-G2 ChIP-seq was made using multiBigwigSummary and plotCorrelation in the deepTools suite (Ramírez et al. 2016).

### Correlation analysis of PfAP2-G2 with other histone marks

Correlations were calculated between the occupancy of PfAP2-G2 and that of various histone marks previously reported in the literature, including H3K9me3, H4ac, H4K20me3, H3K26ac, H3K9ac, H3K4me3 (Karmodiya et al. 2015), H3K36me2 and H3K36me3 (Jiang et al. 2013) and heterochromatin protein 1 (Nicolas M B Brancucci et al. 2014) First, read counts were obtained from PfAP2-G2 and nine histone marks in the gene body of all *P. falciparum* genes (n=5735; PlasmoDB annotations, release 28.0). NCIS (Liang and Keleş 2012) was used to scale a control input experiment to each analyzed signal ChIP-seq experiment. Pairwise Pearson correlation coefficient of normalized read counts across 5735 regions for PfAP2-G2 and nine histone marks. Then, proteins were ordered using complete-linkage hierarchical clustering with Euclidean distance.

### PfAP2-G2-GFP Immunoprecipitation and Mass Spectrometry

Nuclear extraction was performed from the PfAP2-G2::GFP parasites as previously described (Voss et al. 2002). A day before the immunoprecipitation ~1mg of M-270 Epoxy Dynabeads (Invitrogen) per 100 ml of culture were conjugated with polyclonal anti-GFP antibodies at concentration of 5ug/mg (Abcam ab290) or with IgG (Abcam ab46540) overnight at 30°C as previously described (Joshi et al. 2013). Immunoprecipitation was carried out after washing the conjugated beads with dilution buffer and incubating with diluted nuclear extracts for 90 minutes at 4°C. The beads were washed extensively with dilution buffer and 1X PBS and protein was eluted using 20 μl loading buffer by shaking for 10 minutes at 70°C at 1000 rpm and protein was stored at −20°C. Samples were quality controlled by western blotting and subjected to in-gel tryptic digest.

For tryptic digest the samples in polyacrylamide gel slices were destained with 50mM AmBic/50% ACN (Acetonitrile) and dehydrated using 100% ACN. Disulfide bonds in protein were reduced with 10mM DTT and reduced cysteine residues were then alkylated with 50mM Iodoacetamide. Finally, the processed samples were tryptic digested (Trypsin gold, 6ng/μl) overnight at 37°C on a thermomixer, and peptides were extracted from the gels. The dried peptide pellets were analyzed by LC-MS/MS using Q Exactive mass spectrometer (Indiana University Core Proteomics).

The data was processed using Trans-Proteomic Pipeline (TPP) as previously described with a few modifications (Deutsch et al. 2010). Spectra were searched against the *P. falciparum* proteome and contaminant repository database (CRAPome) using tandem and comet searches (Mellacheruvu et al. 2013). This search was combined in InterProphet and the proteins were searched with ProteinProphet. Only proteins with error rate less than 0.01 were reported. The immunopurification assay was performed 3 times and the data was combined to identify the significantly enriched proteins between the IP and control using SAINT (Choi et al. 2011). Proteins with probability score of 0.99 were considered for analysis.

## Supporting information

Singh_PfAP2_G2_Supplementary_Figures

## Additional supplementary materials

**Supplementary Table 1: PfAP2-G2::GFP ChIP-seq data in trophozoite stage** First tab of the table contains all the upstream peaks common in two out of three biological replicates. Second tab has only 86 genes that were considered for the analysis. Indicated are coordinates of each peak, the orientation of the gene, genes downstream to the peaks and distance of peak and ATG of the gene.

**Supplementary Table 2: PfAP2-G2::GFP ChIP-seq data in trophozoite stage** Table contains peaks on the gene-body common in two out of three biological replicates. Also, indicated are coordinates of each peak and genes containing the peaks..

**Supplementary Table 3: PfAP2-G2::GFP ChIP-seq data in gametocyte stage III** Table contains all the upstream peaks common in two biological replicates. Indicated are coordinates of each peak, the orientation of the gene, genes downstream to the peaks and distance of peak and ATG of the gene.

**Supplementary Table 4: PfAP2-G2::GFP ChIP-seq data in gametocyte stage III** Table contains peaks on the gene-body common in two biological replicates. Also, indicated are coordinates of each peak and genes containing the peaks.

**Supplementary Table 5: List of the differentially expressed genes in asexual blood stage** Significance analysis of microarray (SAM) of the asexual stage timecourse data showed that 327 transcripts were differentially expressed (>1.5 log2FC, 0.30 FDR) in the PfAP2-G2 KO line compared to WT. Out of these, 237 showed enhanced transcript abundance in the PfAP2G2 KO line and 90 showed decreased abundance. The first tab of the sheet has upregulated genes and the second tab has downregulated genes along with fold change, q-value and local false discover rate.

**Supplementary Table 6: List of the differentially expressed genes in gametocytes** Significance analysis of microarray (SAM) of the gametocyte timecourse data showed that 444 transcripts were differentially expressed (>1.5 log2FC, 0.30 FDR) in the PfAP2-G2 KO line compared to WT. Out of these, 274 showed enhanced transcript abundance in the PfAP2G2 KO line and 170 showed decreased abundance. The first tab of the sheet has upregulated genes and the second tab has downregulated genes along with fold change, q value and local false discover rate.

**Supplementary Table 7: List of PfAP2-G2 interacting partners** First tab contains the list of peptides obtained in the in the upper band of PfAP2-G2 and the second tab has the list of peptides obtained in the in the lower band of PfAP2-G2 along with the spectral counts, average and maximum probability and FDR. Only the genes with high probability of interaction measured by SAINT (pSAINT 0.9 and 1%FDR) was considered for the analysis.

## Acknowledgements

This project was largely funded by NIH/NIAID 1R01 AI125565 (ML). NY and SM were supported by R01 GM121613 (SM). We thank Kelly Rios for assistance with proteomic data analysis.

