## Supplementary material for "The PfAP2-G2 transcription factor is a critical regulator of gametocyte maturation": Singh_PfAP2_G2_Supplementary_Figures

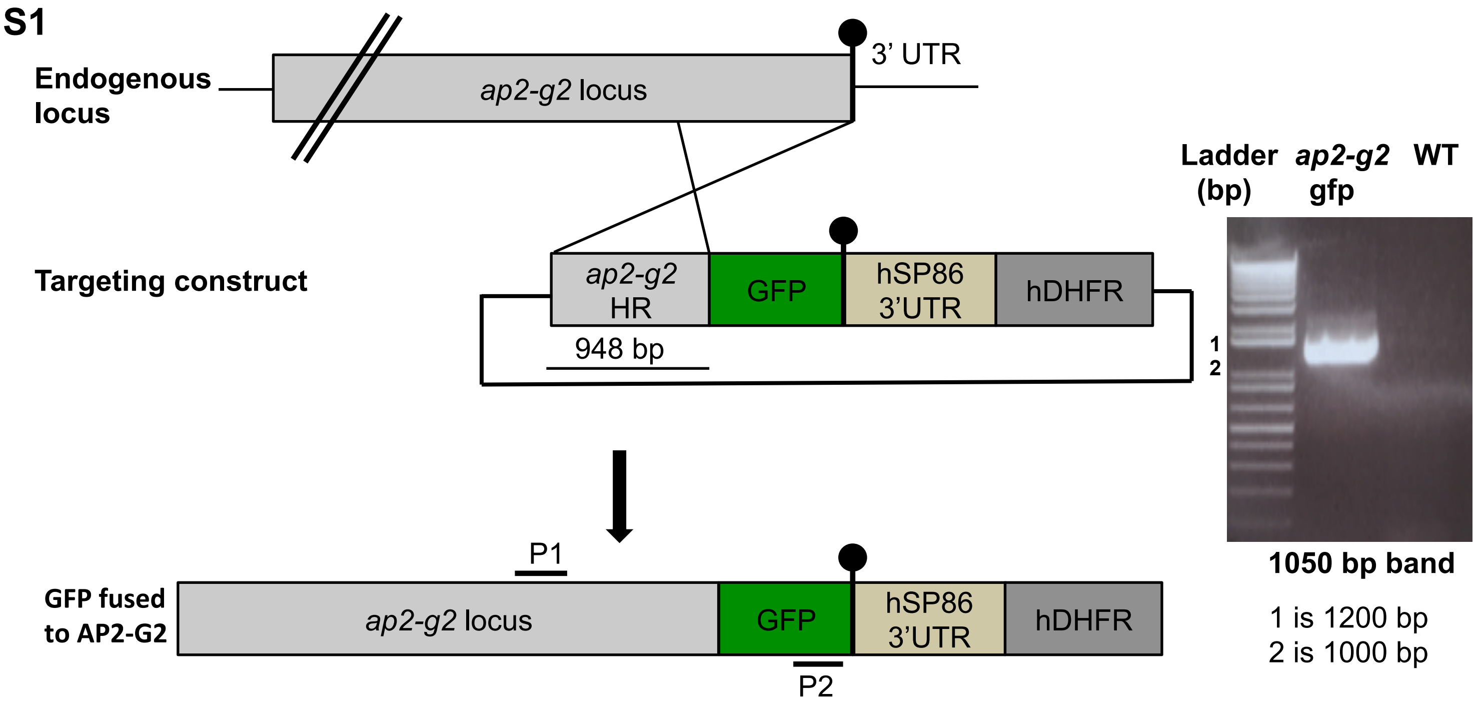


**Sup Fig 1** Schematic showing the strategy used to tag the 3' end of the *pfap2-g2* gene with *gfp*. The tagged parasites were PCR-verified using the primers P1 and P2. A band of the right size was detected only in the transgenic parasite, suggesting the correct modification of the locus.

S2A

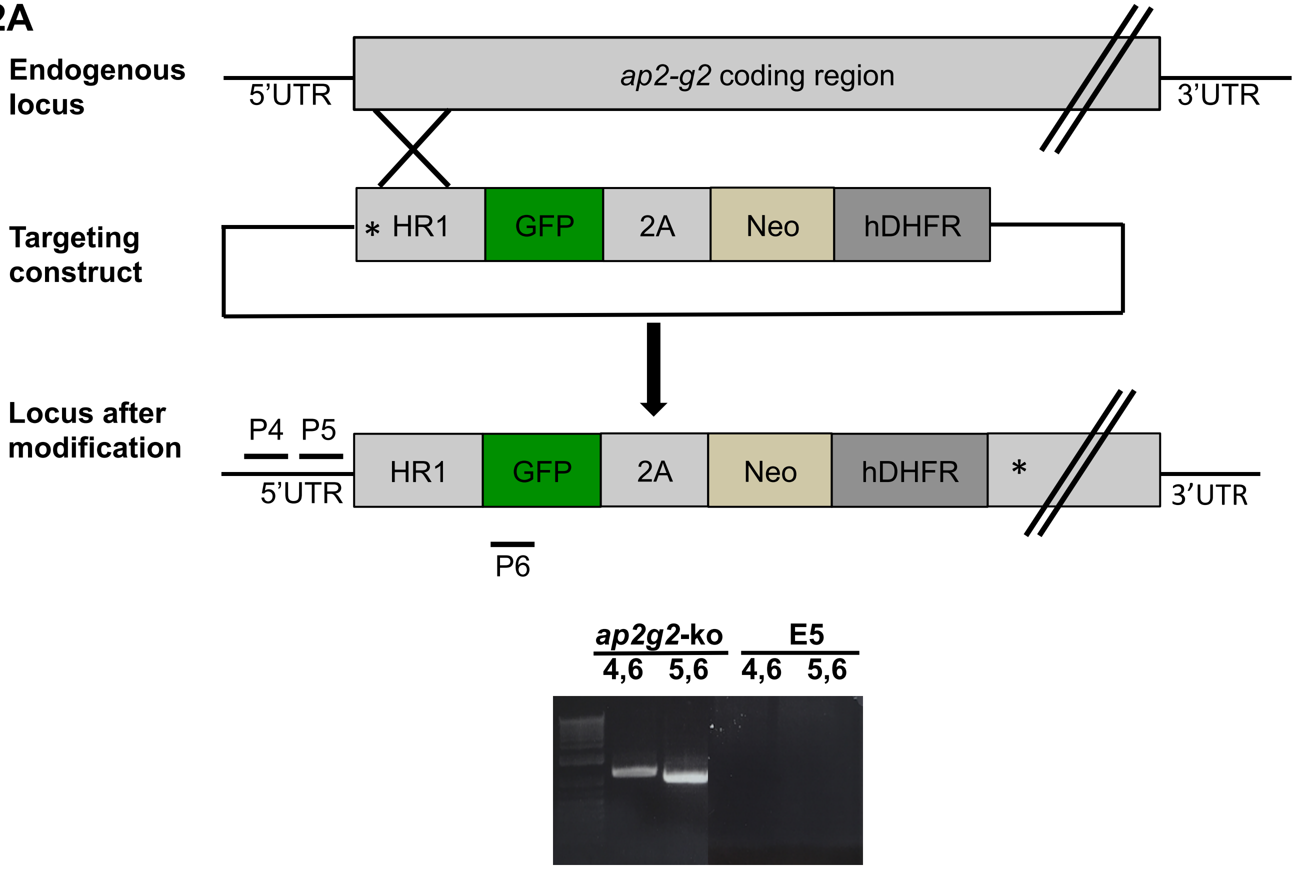


S2B

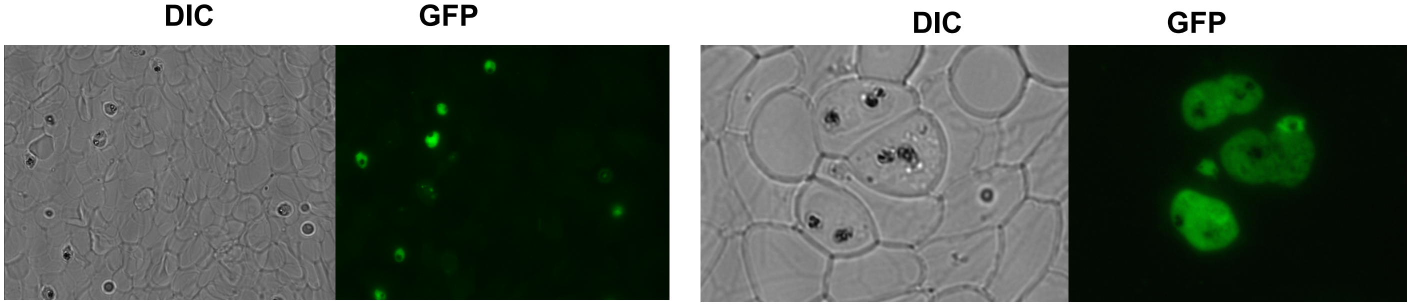


**Sup Fig 2** (A) Schematic showing the strategy used to create the *pfap2-g2* knockout line using the selection linked integration (SLI) method (Birnbaum et al., 2017). A 500 bp homology region with the stop codon (starred) at the beginning ensured that the downstream fragment generated after the single homologous recombination was not expressed. The KO parasite was cloned by limiting dilution and was PCR-verified using the primers P4, P5 and P6. P4 and P5 binds to the 5' UTR upstream of the homology region, whereas primer P6 binds to the GFP, which can generate bands only in recombinant parasites and not in WT. (B) Live fluorescence imaging of the recombinant parasites expressing GFP with a disrupted *ap2-g2* locus in the mixed trophozoite and schizont stages. Image in the left shows that most of the parasites obtained were recombinant and the image in right confirms that stain is diffused throughout the parasites, suggesting that the protein is no longer localized in the nucleus.

**S3**

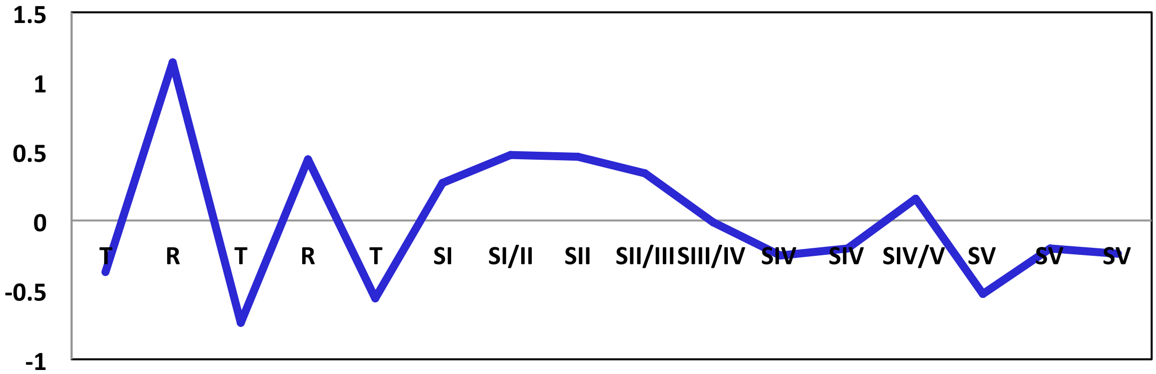


**Sup Fig 3** Graph showing Transcriptional profile of *pfap2-g2* in asexual and gametocyte stages based on the previous research (van Biljon *et al.*, 2019). T is trophozite, R is ring and SI to SV is stage I to stage V of gametocytes.

S4A

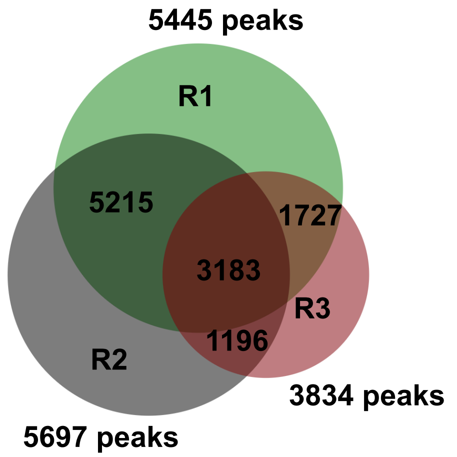


S4B

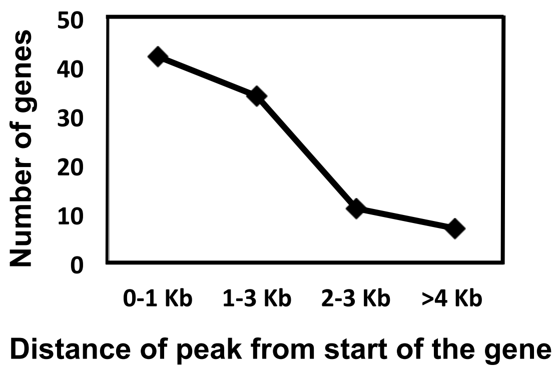


S4C

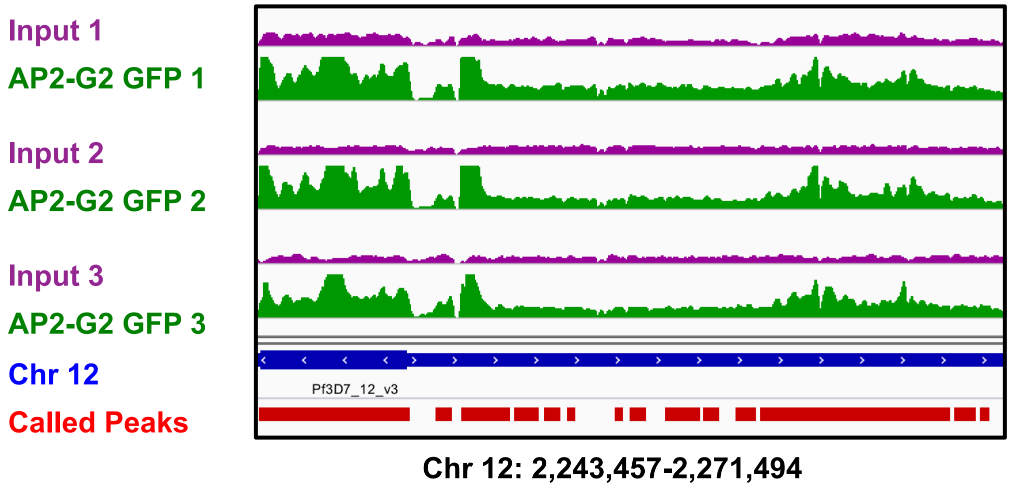


**Sup Fig 4** (A) Distance of peaks from the start of the gene. Out of 119 genes with upstream PfAP2-G2 peaks, ~100 has peaks located within 3Kb from the ATG of the gene. (B) Regions occupied by PfAP2-G2 at the end of chromosomes in all three replicates. Thick blue tracks represent the chromosome and red tracks represents the called peaks by MACS2**.** (C) Enrichment of PfAP2-G2 at the end of chromosomes in all three replicates.

S5

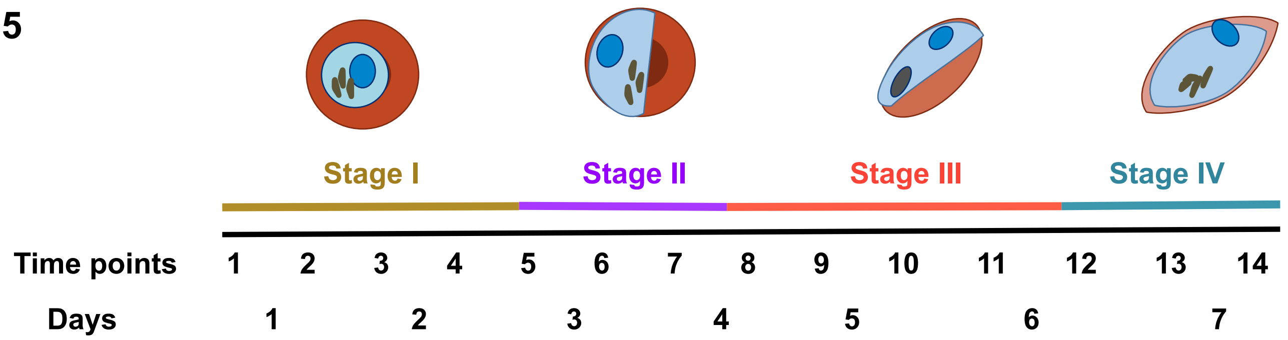


**Sup Fig 5** Schematic showing mRNA collection time-points for gametocyte time course.Gametocyte induction was performed when the ring parasitemia was around 7-10%. The next cycle rings were set as day 1 of gametocytogenesis as some are sexually committed rings. To prevent reinvasion in the next cycle and thus formation of new asexuals, parasites were treated with 20U/ml heparin from day 1 post-induction through day 4. RNA was collected every 12 hours from day 1 to day 14 (stage I to stage IV).

S6

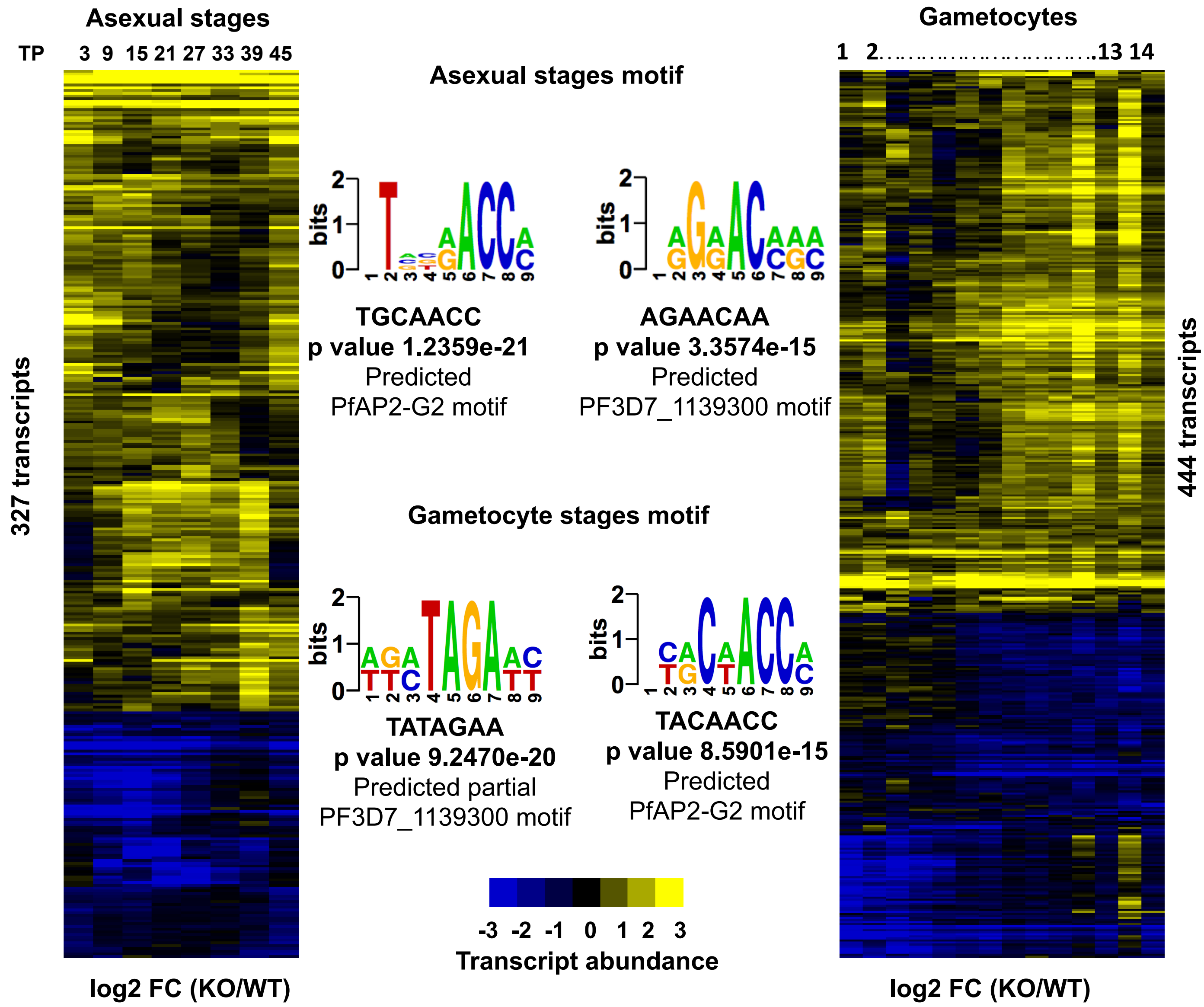


**Sup Fig 6** Shown here are the heatmaps of the significantly changing transcripts in the PfAP2-G2 KO line compared to WT (>1.5 log2FC, 0.30 FDR) for both asexuals and gametocytes. Also, shown are the DNA sequence motif associated with the 5' upstream region of (~1000bp from the ATG) for genes with increased transcript abundance in asexual stage using finding informative regulatory elements (FIRE) algorithm. We also recover similar motif from the genes with increased and decreased transcript abundance in gametocytes.

**S7**

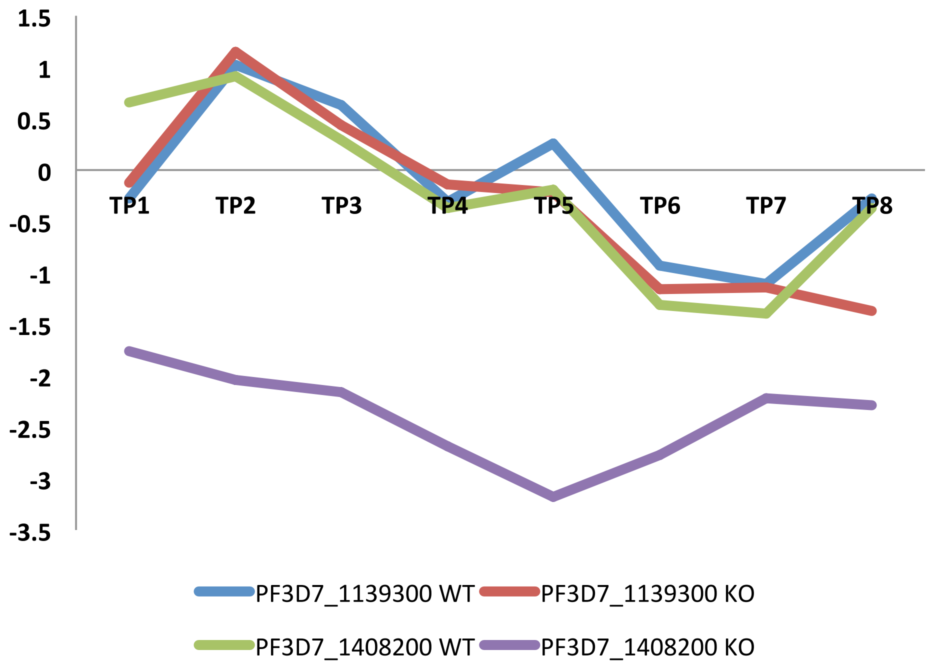


**Sup Fig 7** Plot showing mRNA Expression level of PfAP2-G2 (PF3D7_1408200) and PF3D7_1139300 suggests that both are co-expressed.

**S8**

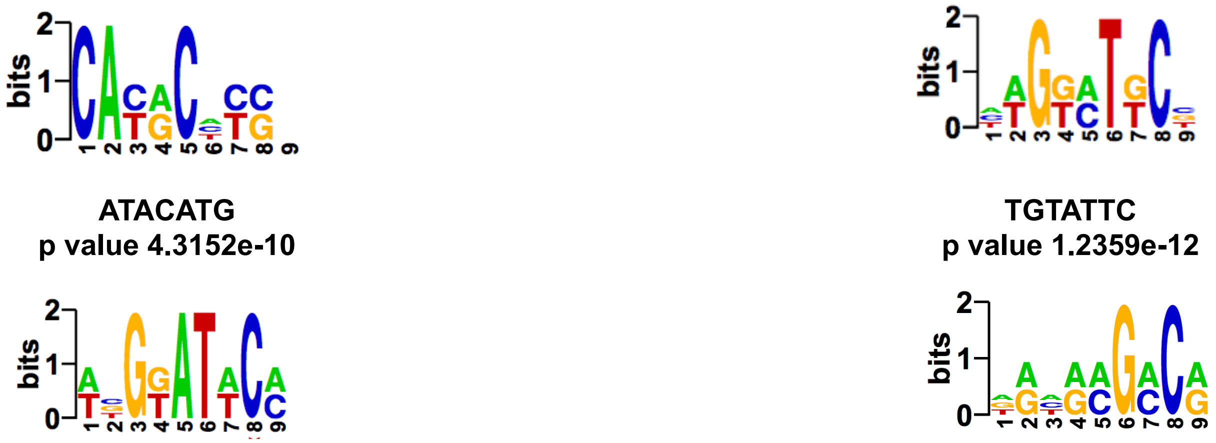

**ATACATG**
**p value 4.3152e-10**

**TGTATTC**
**p value 1.2359e-12**

**TGTATTC**
**p value 4.5709e-08**

**AAAAGAC**
**p value 4.4668e-12**
Predicted partial
Motif associated with
gametocyte genes

**Sup Fig 8** Shown are the DNA sequence motif associated with the 5' upstream region of (~1000bp from the ATG) for genes with decreased transcript abundance in asexual and gametocyte stages using finding informative regulatory elements (FIRE) algorithm.

**S9**

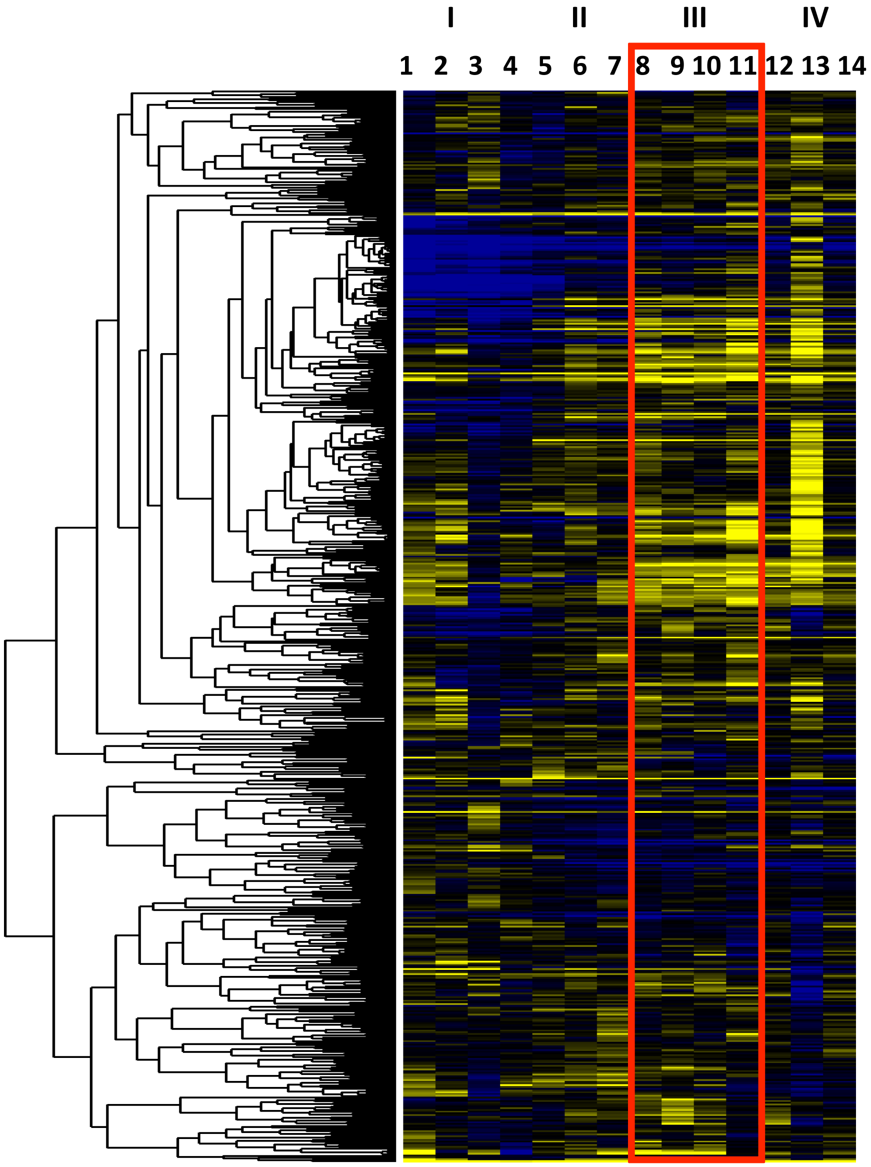


**Sup Fig 9** Expression fold change (log2 KO versus WT) for the genes that are bound by PfAP2-G2 in the in the gene body in the gametocyte stage III. The genes are clustered based on Pearson’s correlation. Boxed is the timepoint at which ChIP-seq was performed.
